# GBScleanR: Robust genotyping error correction using hidden Markov model with error pattern recognition

**DOI:** 10.1101/2022.03.18.484886

**Authors:** Tomoyuki Furuta, Toshio Yamamoto, Motoyuki Ashikari

## Abstract

Reduced-representation sequencing (RRS) provides cost-effective and time-saving genotyping platforms. Although the outstanding advantage of RRS in throughput, the obtained genotype data usually contains a large number of errors. Several error correction methods employing hidden Morkov model (HMM) have been developed to overcome these issues. Those methods assume that markers have a uniform error rate with no bias in the allele read ratio. However, bias does occur because of uneven amplification of genomic fragments and read mismapping. In this paper we introduce an error correction tool, GBScleanR, which enables robust and precise error correction for noisy RRS-based genotype data by incorporating marker-specific error rates into the HMM. The results indicate that GBScleanR improves the accuracy by more than 25 percentage points at maximum as compared to the existing tools in simulation datasets and achieves the most reliable genotype estimation in real data even with error prone markers.

## Introduction

The rise of next generation sequencing (NGS) has opened the door to a new generation of genetic and genomic research^1^. NGS provides genotyping platforms with dramatically improved flexibility and throughput that are now routinely applied to diverse fields of biology in which information about the nucleotide sequences of organisms is paramount^2–4^. The cost of NGS has decreased significantly over the past decade. Nevertheless, obtaining a genotype via whole genome resequencing is still an expensive option for researchers, particularly those working on breeding projects that involve the genomes of thousands of individuals^5^. Many different RRS-based genotyping methods have been introduced to meet the demands for cost-effective genotyping systems with dense markers ^6^. RRS is a sequencing technique using NGS with reducing sequence targets by taking reads only from the limited portion of a genome^6^. Since restriction-site associated DNA sequencing (RAD-seq) was first published in 2008, several derivative methods have been introduced, including methods such as ddRADseq, nextRAD, ezRAD, 2b-RAD, and RAD Capture^7–12^. Genotyping-BY-Sequencing (GBS) and its derivative, two-enzyme GBS, are also outstanding RRS methods^13,14^. GBS has already been widely applied to many crops and animals, such as rice, sorghum, chickpea, cattle, and mice^15–19^. GBS, RAD-seq and its derivatives basically obtain sequences from a limited portion of a genome by fragmentating genomic DNA with restriction enzymes before sequencing from adapters that are ligated at the cut site. This reduction in the number of sequence targets allows for multiplexed sequencing, freeing researchers from the laborious pre-surveys that are usually required to find useful genotyping markers for PCR-based and hybridization-based genotyping systems such as simple sequence repeat (SSR) genotyping and SNP genotyping array^20,21^. However, RRS has well-known drawbacks that are derived from its technological nature and include mismapped reads and the undercalling of heterozygous genotypes. Although the most significant advantage of RRS lies in its multiplexibility, this characteristic reduces the number of reads per NGS run. Since the sequences that can be retrieved via NGS are highly stochastic, the limited number of reads can result in the undercalling of heterozygotes, in which recovering only one allele at a heterozygous site leads to its incorrect identification as a homozygote. The lack of reads, in addition, also generate a substantial number of missing genotype calls.

Several error correction tools have been developed to overcome these disadvantages, with two categories available, one of which imputes the missing genotype calls and the other both imputes the missing genotype calls and corrects the undercallings. Beagle is the most popular example of the former, and is a tool for the haplotype-phasing that is frequently used in genome-wide association study (GWAS) that can infer missing genotypes based on the phased haplotype information^22^. The method that is used to correct the undercalling of heterozygotes was first published in 2009, and determines a genotype from the allele read ratio within a sliding window^23^. Recent publications have introduced more statistically sophisticated methods that use HMMs, such as FSFHap, TIGER, LB-Impute, and magicImpute^24–27^. An HMM is a statistical model in which it is assumed that a series of observed data is the output of a sequence of hidden states that follow a Markov process^28^. In the case of genotyping data, the series of observed data and the sequence of hidden states correspond to the observed genotype calls (or allele read counts) and the true genotype along a chromosome, respectively. The existing error correction methods provide a robust estimation of the true genotype against a background that includes the undercalling of heterozygotes and missing calls that result from stochastic error.

The existing error correction methods, however, assume constant error rates for all markers. For example, those assume a 50:50 probability that a read could be obtained for one of two possible alleles at a heterozygous site. Although this assumption is true in an ideal situation, in practice, genotype data contains a significant number of error prone markers that show skewed probabilities in the allele read acquisition as a result of actual biological unevenness^29–31^. Polymorphisms that exist between the genomes of samples change the fragmentation patterns of the different genomes. Even if two independent reads are mapped at the same locus, the sequences of the genomic fragments from which each of the two reads originated can vary in terms of GC content and length, and sometimes include large insertions or deletions in the unsequenced region of the fragments. Since the GC contents and the lengths of the restricted fragments are known to affect the amplification efficiency, the probability of observing a read for either allele may differ. These biases are likely to be more prominent in a polymorphism-rich population, for example, that derived from a cross between distant relatives. Hence, mismatches occur between the real data, which shows marker specific error rates, and the models that assume the uniform error rate. Those mismatches then result in biased genotype estimation and poor error correction accuracy.

Here we introduce an R package “GBScleanR” that implements an HMM-based error correction algorithm. Our algorithm estimates the allele read bias and mismap rate per marker and incorporates those into the HMM as parameters to capture the skewed probabilities in read acquisitions. This paper demonstrates a comparison of GBScleanR and two well established error correction tools; LB-Impute and magicImpute^26,27^. While LB-Impute accepts only biparental populations that are derived from inbred founders, magicImpute can work on bi- and multi-parental populations that are derived from both inbred and outbred founders. The magicImpute algorithm is also able to estimate founder genotypes simultaneously with the offspring genotypes, meaning that it does not require complete and high-quality information concerning founder genotype. Similar with magicImpute, GBScleanR also supports many scenarios in which experimental crosses are usually conducted and is designed to estimate founder and offspring genotypes simultaneously with higher robustness for error prone markers. We first show simulation studies to present the accuracy and robustness of GBScleanR using simulation datasets that have severe allele read biases at error prone markers. The simulation assumes three scenarios; a biparental F2 population (homoP2_F2), an outbred F2 population (hetP2_F2), and an 8-way recombinant inbred line (homoP8_RIL). We then demonstrate the reliability and robustness of GBScleanR using the real data derived from a cross between distant relatives of rice, which potentially contains a number of error prone markers.

## Results

### Evaluation using simulation datasets

We first evaluated the three algorithms using the simulation datasets in the scenarios homoP2_F2, hetP2_F2, and homoP8_RIL. If no allele read bias was observed, the proportion of reference allele reads per marker would be 0.5 on average. However, the values that were derived from the real data showed a highly biased distribution, with a large number of markers showing bias towards more reference allele reads than expected (Supplemental Figure S1). Some markers also showed more alternative than reference allele reads. The simulation datasets mimicking allele read biases observed in the real data were subjected to error correction via GBScleanR, magicImpute, and LB-Impute. The superiority of GBScleanR lies in the iterative parameter optimization (IPO) in which allele read biases and mismap rates of markers are iteratively estimated. Thus, we also tested GBScleanR without IPO to demonstrate the effect of IPO on genotype estimation.

Under the condition of such biases, the genotype estimations made by the algorithms was highly disturbed, except for GBScleanR with IPO (Figure 1 and Supplemental Data S1). In the homoP2_F2 scenario, GBScleanR with IPO scored the best correct genotype call rates for almost all conditions (Figure 1A-C). When the simulation datasets were generated without allowance to have no read at some markers in founders (indicated as “nonzero” in Figure 1), LB-Impute showed the highest correct call rates for the datasets with a 0.1× offspring read depth. However, LB-Impute also scored the highest miscall rate for those datasets (Supplemental Data 1). The reduction of accuracy was observed in all of the tested algorithms, when missing genotypes due to no read observation at some markers were allowed for founders (shown as “allowzero” in Figure 1). Nevertheless, GBScleanR was less affected by missing genotypes in founders and outperformed the other algorithms. In addition, GBScleanR with IPO only kept increasing the estimation accuracy as offspring read depth increased, while the other algorithms showed a plateau at 0.5-20x in the results for the “nonzero” datasets (Figure 1A-C). The difference in correct call rates between GBScleanR with IPO and the other algorithms reached more than 25 percentage points at maximum in the datasets with a 20× offspring read depth. On the other hand, the algorithms except for GBScleanR with IPO showed an increase of the miscall rate as the read depth increased (Supplemental Figure 2).

**Figure 1.**
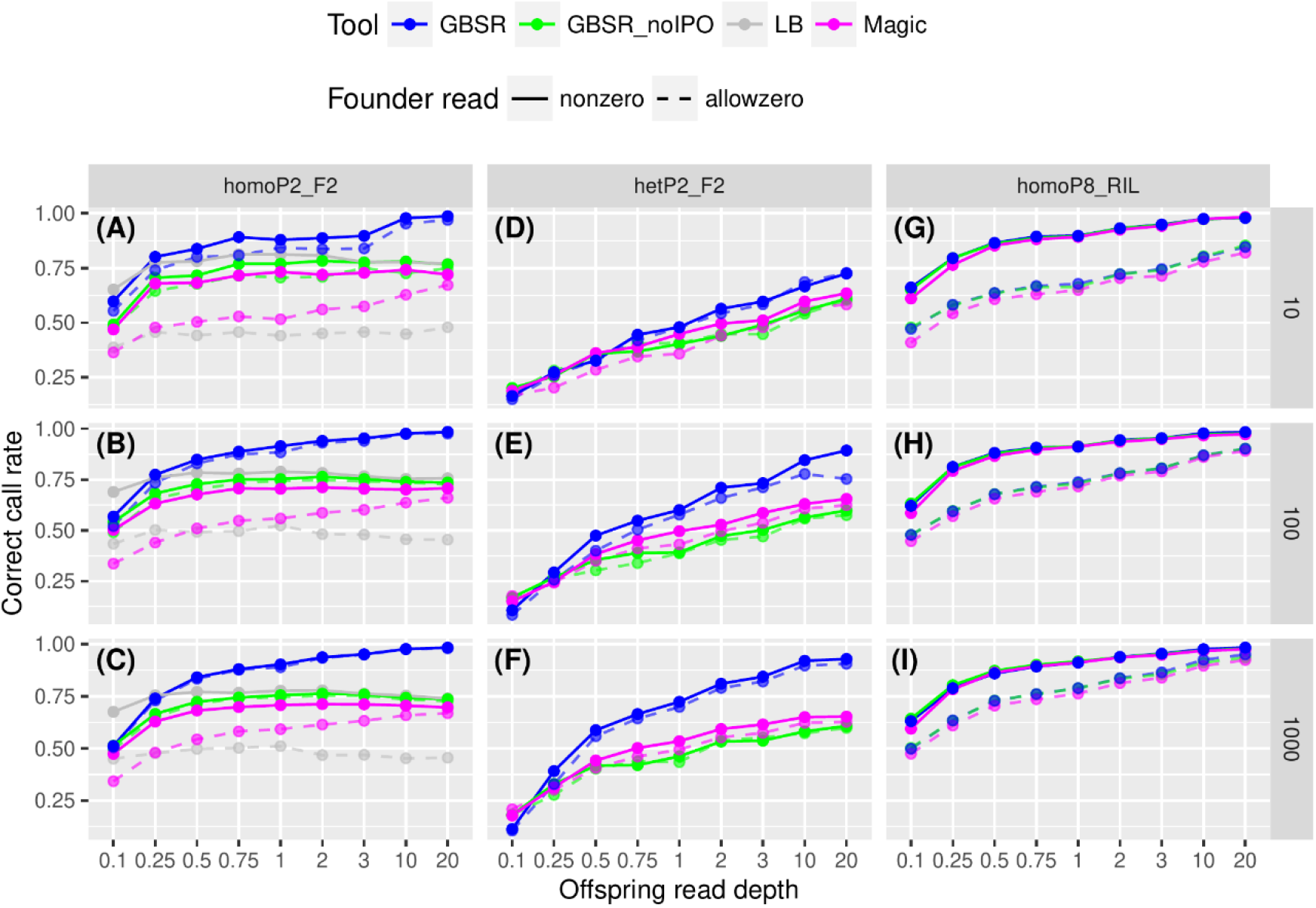
Offspring genotype estimation accuracies for the simulation data. The plots show correct call rates for the datasets with given read depths for offspring (x axis) in the homoP2_F2 (A-C), hetP2_F2 (D-F), and homoP8_RIL (G-I) scenarios. The rows of the panels indicate the differences in the number of samples in the simulated data as shown in the strips on the right. Solid lines and dashed lines represent the results for the datasets with (allowzero) or without (nonzero) missing of founder reads. “GBSR” and “GBSR_nolPO” represent GBScleanR with and without IPO. “LB” and “Magic” indicate LB-Impute and magicImpute, respectively.

Similar with the results in homoP2_F2, GBScleanR showed the best correct call rates in almost all datasets of hetP2_F2 and the difference in the scores between the algorithms reached more than 25 percentage points at maximum (Figure 1D-F). Although the accuracy increased as the number of samples and reads increased, the lower absolute scores were observed than those observed in the datasets of homoP2_F2, indicating the difficulty of the genotype estimation on a population derived from outbred founders having heterozygous genomes. Unlike the homoP2_F2 datasets, the genotype estimation accuracy for the hetP2_F2 datasets was highly affected by the number of samples (Figure 1A-F). The increase in correct call rate between the datasets with 10 and 1000 samples reached around 26 percentage points at maximum in the hetP2_F2 datasets with a 0.5× offspring read depth, whereas the maximum increase of the score in the homoP2_F2 datasets was about only 5 percentage points (Supplemental Data 1).

Unlike the F2 datasets, we could not find any remarkable difference in the accuracy of genotype estimation for the homoP8_RIL datasets (Figure 1G-I). This result seems to have reflected the higher homozygosity in the simulated genotypes. Since the offspring in the homoP8_RIL datasets had homozygous genotypes at almost all markers due to repeated self-fertilization, only the reads of either allele representing the true genotype could be observed even at error prone markers with allele read biases.

### Estimation of founder genotype and phased genotype

GBScleanR and magicImpute have been designed to simultaneously estimate the genotypes of both founders and offspring. Thus, we next compared the founder genotype estimation accuracy of the algorithms. As expected from the estimation accuracy for offspring, GBScleanR outperformed magicImpute (Figure 2 and Supplemental Data 2). While both algorithms showed similar and high correct call rates in the homoP2_F2 and homoP8_RIL datasets when “nonezero”, magicImpute resulted in less accurate founder genotype estimations compared with GBScleanR for the “allowzero” datasets (Figure 2A-C and G-I). Unlike the offspring genotype estimation, IPO did not affect the founder genotype estimation in the homoP2_F2 and the homoP8_RIL datasets (Figure 2A-C and G-I). Improvement in the estimation accuracy by IPO was observed only in the hetP2_F2 datasets with a relatively large number of offspring and offspring reads (Figure 2D-F). This might be because heterozygotes in the founder genotypes were only assumed in the hetP2_F2 datasets but not in the others; the homozygous genotypes of the inbred founders were not affected by allele read biases.

**Figure 2.**
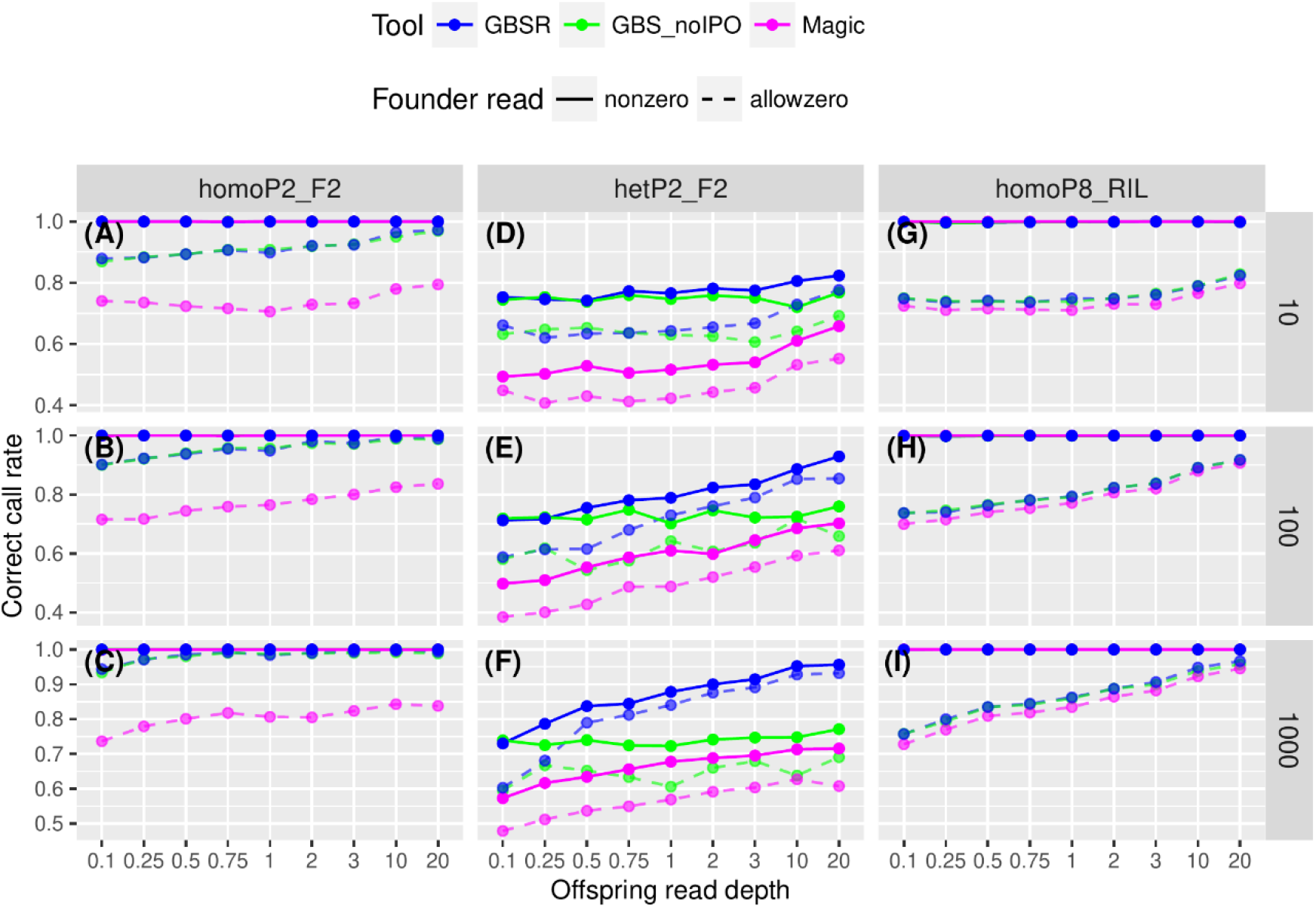
Founder genotype estimation accuracies for the simulation data. The plots show correct call rates for the datasets with given read depths for offspring (x axis) in the homoP2_F2 (A-C), hetP2_F2 (D-F), and homoP8_RIL (G-I) scenarios. The rows of the panels indicate the differences in the number of samples in the simulated data as shown in the strips on the right. Solid lines and dashed lines represent the results for the datasets with (allowzero) or without (nonzero) missing of founder reads. “GBSR” and “GBSR_noIPO” represent GBScleanR with and without IPO. “LB” and “Magic” indicate LB-Impute and magicImpute, respectively.

In addition to the genotype information of offspring and founders, both GBScleanR and magicImpute provide phasing information for the offspring genotypes. Phased genotype, which is also referred to as haplotype, is the sequence of alleles on each of a chromosome pair. Correct call rates were evaluated by scoring the proportion of estimated haplotype that matched with the simulated haplotype. As expected from the superiority for both founder and offspring genotype estimation, GBScleanR also outperformed magicImpute in the haplotype estimation (Supplemental Figure 3 and Supplemental Data 3). Although allowance of no read in founders appears to have not remarkably affected the accuracy in most cases, the “nonzero” datasets resulted in better estimations than those of “allowzero” in the homoP2_F2 datasets, especially in the results of magicImpute.

### Less biased estimation by GBScleanR

We further evaluated the quality of the estimated genotype data by visualizing the estimated genotype ratio at each marker (Figure 3 and Supplemental Figure 4 and 5). This visual inspection employed the homoP2_F2 datasets that have 1000 offspring with offspring reads depths of 0.5×, 3×, and 20× without allowance of no read in founders. In an F2 population derived from inbred parents, we can expect a 1:2:1 genotype ratio in offspring if no segregation distortion. As shown in Figure 3, the estimations by GBScleanR showed 1:2:1 segregation at almost all markers, which nearly completely overlapped with the true genotype ratio throughout the chromosome, except for the dataset with only 0.5× offspring read (Figure 3A-C). In addition, miscalls in the estimated genotype data were distributed almost uniformly throughout the entire chromosome with very small amount per marker. In contrast, LB-Impute showed severely biased genotype ratios with high miscall rates clustered at several loci on the chromosome (Figure 3D-F). The increase of offspring read depth made biased genotype estimation severer and increased miscall rates as also shown in Supplemental Figure 2. Similar results were observed in the genotypes estimated by magicImpute (Figure 3G-I). Even though less miscalls were generated in magicImpute compared to LB-Impute, many regions showed the biased estimation of reference homozygotes, in which reference homozygotes were preferentially estimated while heterozygotes were left as missing, particularly visible in the dataset with 0.5× offspring read depth (Figure 3G). The loci showing higher miscall rates overlapped in the plots for LB-Impute and magicImpute. Thus, these miscalls and biased genotype estimations seemed to be caused by error prone markers at which severe allele read biases prevent the algorithms from correct genotype estimation. Similar results were observed in the datasets with 10 and 100 offspring (Supplemental Figure 4 and 5). Although the true genotype did not show 1:2:1 ratio due to the small population size in the datasets with 10 offspring, GBScleanR showed the estimated genotype ratio that highly overlapped with the true ratio (Supplemental Figure 4A-C). These results demonstrated the robustness of genotype estimation by GBScleanR even at error prone markers.

**Figure 3.**
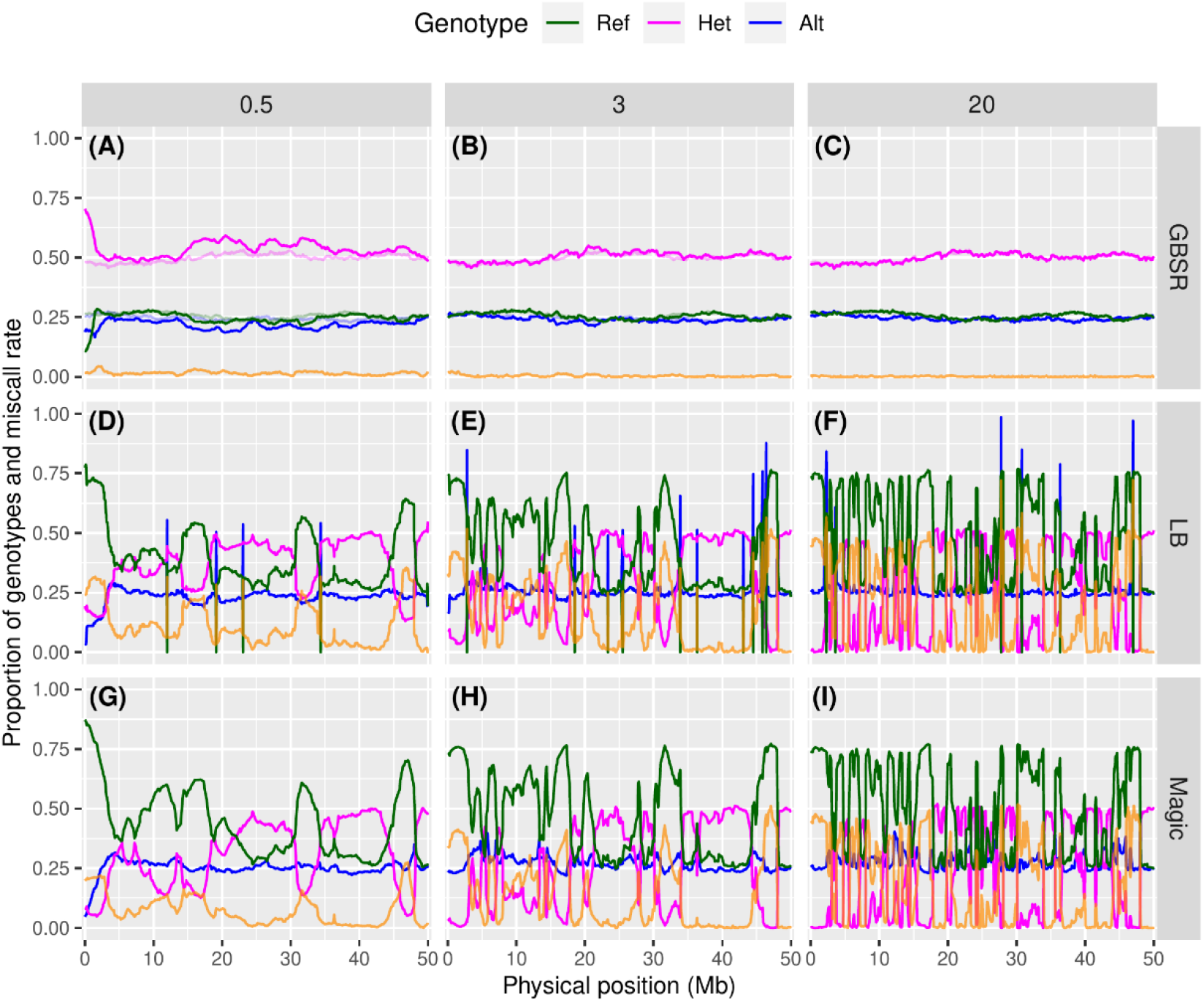
Genotype ratios at markers ordered along a simulated chromosome. The plots show genotype ratios calculated from the estimated genotype data for the homoP2_F2 dataset with 1000 offspring and 620 markers with offspring read depths of 0.5×, 3×, and 20× and without allowance of no read in founders. “GBSR” (A-C), “LB” (D-F), and “Magic” (G-I) indicates estimated genotype by GBScleanR, LB-Impute, and magicImpute, respectively. Proportions of reference homozygous, heterozygous, and alternative homozygous genotypes are represented by green, magenta, and blue lines, respectively. Orange lines indicate miscall rates at markers. True genotype ratios are indicated by transparent lines in the panels for GBScleanR (A-C).

### Evaluation using real data

We finally evaluated the algorithms using the real data that was the genotype data obtained by GBS on an F_2_ population consist of 814 individuals derived from a distant cross between *Oryza sativa* and *O. longistaminata*. As usually conducted to evaluate the performances of genotpye error correction tools for real data, the concordance rate of the estimated genotype calls was measured using the masked genotype data derived from the real data. First of all, GBScleanR scored the best concordance rate in all chromosomes except for chromosome 11 in the comparison with LB-Impute and magicImpute (Figure 4A and Supplemental Data S4). Relatively large differences were observed in chromosome 7, 8, and 10, in which GBSclenaR scored 10.5, 12.5, and 13.0 percentage points higher concordance rates than magicImpute. On the other hand, chromosome 5 showed only a tiny difference in the concordance rate that was 1.5 percentage points between GBScleanR and magicImpute. The chromosomes with severer allele read biases, e.g. chromosome 7-12, seemed to tend to show smaller concordance rates than those observed in the other chromosomes (Figure 4A and Supplemental Figure 1).

**Figure 4.**
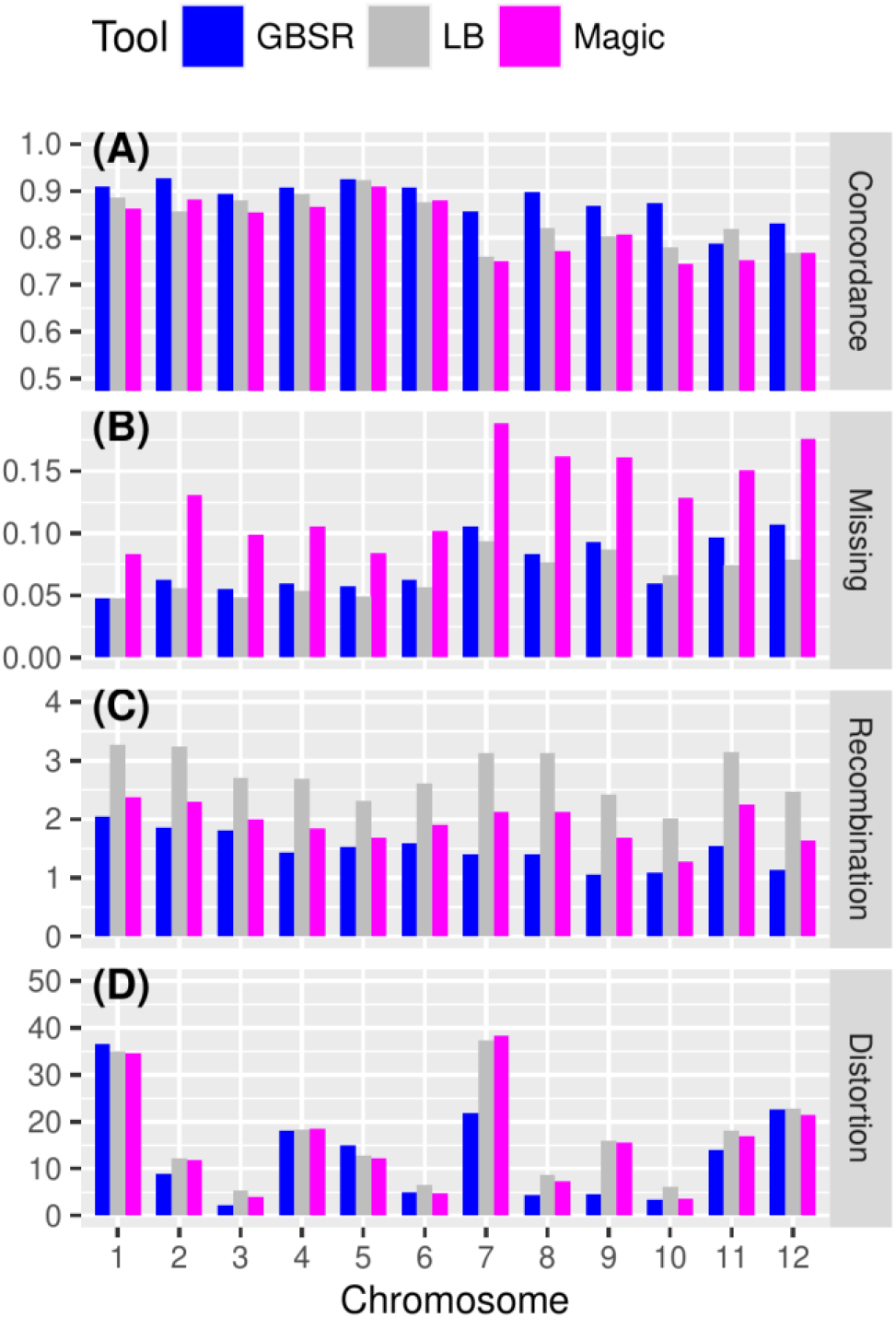
Evaluation of genotype estimation accuracy using the real data. (A) Concordance rate of true genotype calls and estimated genotype calls as the masked genotype calls are shown for each chromosome. The averages of missing rate (B), the number of recombinations (C), and segregation distortion level (D) are also visualized as bar plots for each chromosome. Blue, gray, and magenta bars indicate the values of GBScleanR (GBSR), LB-Impute (LB), and magicImpute (Magic), respectively

Even though GBScleanR showed better concordance rates than the others, the genotype concordance does not directly mean the higher accuracy because we cannot know the true genotype of the real data and there is no evidence whether the genotype calls selected for masking were truly reliable or contained a large number of erroneous genotype calls. To further evaluate the performance of the algorithms, we then calculated the averages of missing rates and the number of recombinations in each chromosome. In the case of missing rate, the highest values were scored by magicImpute for all chromosomes, while GBScleanR and LB-impute showed the similar scores in a majority of chromosomes (Figure 4B and Supplemental Data 4). Similar with the case of the concordance rate, the chromosomes having severer allele read biases showed relatively higher missing rates. A clear difference between the algorithms was observed in the number of recombinations (Figure 4C and Supplemental Data 4). LB-Impute always estimated the most frequent recombinations followed by magicImpute, whereas GBScleanR showed the fewest numbers of recombinations in all chromosomes. The previous study reported that the shortest and longest chromosomes in the estimated genetic map of rice are chromosome 10 and chromosome 1, which are 83.8 and 181.8 cM, respectively^32^. Therefore, the expected numbers of recombinations per chromosome would range 1.6-3.6. The results of LB-Impute seemed to fit to the expected values more than those of GBScleanR. To confirm whether LB-Impute truly estimated more accurate numbers of recombination events, we measured the genomic segment lengths between the recombination breakpoints in chromosomes (Table 1 and Supplemental Data 5). We found that LB-Impute estimated much more numbers of short segments. The genotype data estimated by GBScleanR had 261 and 668 segments in the ranges of 0-1 and 1-2 Mb, respectively (Table 1). On the other hand, LB-Impute estimated the genotype data with 2164 and 4166 of 0-1 and 1-2 Mb segments, respectively. The magicImpute algorithm generated a fewer number of short segments than LB-Impute but a larger number than GBScleanR, which were 627 and 1716. In addition, GBScleanR, magicImpute, and LB-Impute estimated 58, 235, and 948 of 0-1 Mb segments that were heterozygous and flanked by the same homozygous genotypes at the both sides, which indicate double crossovers within 1 Mb stretches (Table 1). The probability of a double crossover within 1 Mb is approximately 0.08% when 1 Mb equals to 4 cM. Thus, the expected number of double crossovers is less than 2 in the population with 814 individuals derived from 1628 gametes. Even though the 58 double crossovers within 1Mb stretches generated by GBScleanR was too much frequent for an ideal F_2_ population, considering the distant cross to generate the given population, these results suggested that GBScleanR estimated more probable and acceptable numbers of recombinations. On the other hand, unexpectedly large numbers of short segments were estimated by LB-Impute and magicImpute, which indicated double crossovers at approximately 4- and 16-fold higher frequencies than those estimated by GBScleanR. Those excess crossovers in short distances seemed to push up the numbers of recombinations artificially.

**Table 1.** The number of genome segments.

|  |  | Segment length |  |
| --- | --- | --- | --- |
|  |  | 0-1 Mb | 1-2 Mb |
| Total<sup>a</sup> | GBScleanR | 261 | 668 |
|  | magicImpute | 627 | 1716 |
|  | LB-Impute | 2164 | 4166 |
| Double<br>crossovers<sup>b</sup> | GBScleanR | 58 | 78 |
|  | magicImpute | 235 | 558 |
|  | LB-Impute | 948 | 1395 |

<sup>a</sup> The total number of genome segments that have a same genotype continuously with the length of 0-1 or 1-2 Mb.
<sup>b</sup> The number of genome segments that have a heterozygous genotype continuously with the length of 0-1 or 1-2 Mb, and flanked by homozygous segments of a same allele.

At last, we measured the segregation distortion level in the estimated genotype data (Figure 4D and Supplemental Data 4). Since our real data was obtained from the rice F_2_ population, it could be expected that any markers show a genotype segregation ratio of 1:2:1 for reference homozygotes, heterozygotes, and alternative homozygotes although segregation distortion due to the reproductive isolation could occur. Relatively low distortion level was observed in the genotype data estimated via GBScleanR compared with those by LB-Impute and magicImpute (Figure 4D). Chromosome 1 was the most distorted one in the genotype data estimated by GBScleanR, while the other algorithms also showed similar distortion levels in this chromosome. On the other hand, the highest distortion level was observed in chromosome 7 in the cases of LB-Impute and magicImpute (Figure 4D). Unlike chromosome 1, GBScleanR showed a much smaller distortion level that was about a half of the levels scored by the other algorithms. In contrast, the three algorithms all resulted in the mildest distortion in chromosome 3 (Figure 4D). To visually inspect the segregation distortion levels, we created the line plots of the estimated genotype ratios (Figure 5 and Supplemental Figure 6). Chromosomes 1, 3, and 7 were selected to demonstrate the negative impact of allele read biases (Figure 5). The line plots indicated that all of the algorithms estimated quite similar genotypes for chromosome 1 and 3, while clearer differences were observed in the plots for chromosome 7. As fewer markers in chromosome 1 and 3 showed sever allele read biases, while chromosome 7 showed relatively higher number of markers preferentially accumulated reference allele reads, chromosome 1 and 3 might have less error prone markers (Supplemental Figure 2). These chromosomes with relatively less error prone do not seem to have made a sever negative impact on the estimation even by LB-Impute and magicImpute, and resulted in the similar estimation with GBScleanR (Figure 5A-F). Since all algorithms resulted in similar distortion patterns around 30 Mb of chromosome 1, the distortion on chromosome 1 was not likely to be due to the misestimation but the true genetic characteristic of this population (Figure 5A-C). On the other hand, slight distortions at 5, 10, 35 Mb of chromosome 3 that were observed in the genotypes estimated by LB-Impute and magicImpute, but not by GBScleanR, might be artifacts produced by biased estimation at error prone markers (Figure 5D-F). In the case of chromosome 7, however, LB-Impute and magiImpute obviously failed in estimating the genotypes and produced artificially distorted genotype ratio patterns (Figure 5G-I). In contrast, GBScleanR showed a smooth and less fluctuating change of the genotype ratio, which was an acceptable pattern as the genotype ratio of a F2 population, throughout the chromosome although severe distortion was observed probably due to true segregation distortion (Figure 6I). The other chromosomes also exhibited similar results to those observed for chromosomes 1, 3, and 7 (Supplemental Figure S14). LB-Impute and magicImpute always resulted in highly biased genotype estimation at many loci and fluctuated genotype ratio throughout the chromosomes. In clear contrast to those results, GBScleanR showed relatively smooth and gradual change of genotype ratio throughout the chromosomes. Even though some loci showed distortion of genotype ratio, majority of chromosomal region had ratio relatively near to 1:2:1 in almost all chromosomes in the result of GBScleanR than those of the other algorithms. Overall, the performance evaluation on the real data demonstrated the superiority and robustness of GBScleanR against error prone markers.

**Figure 5.**
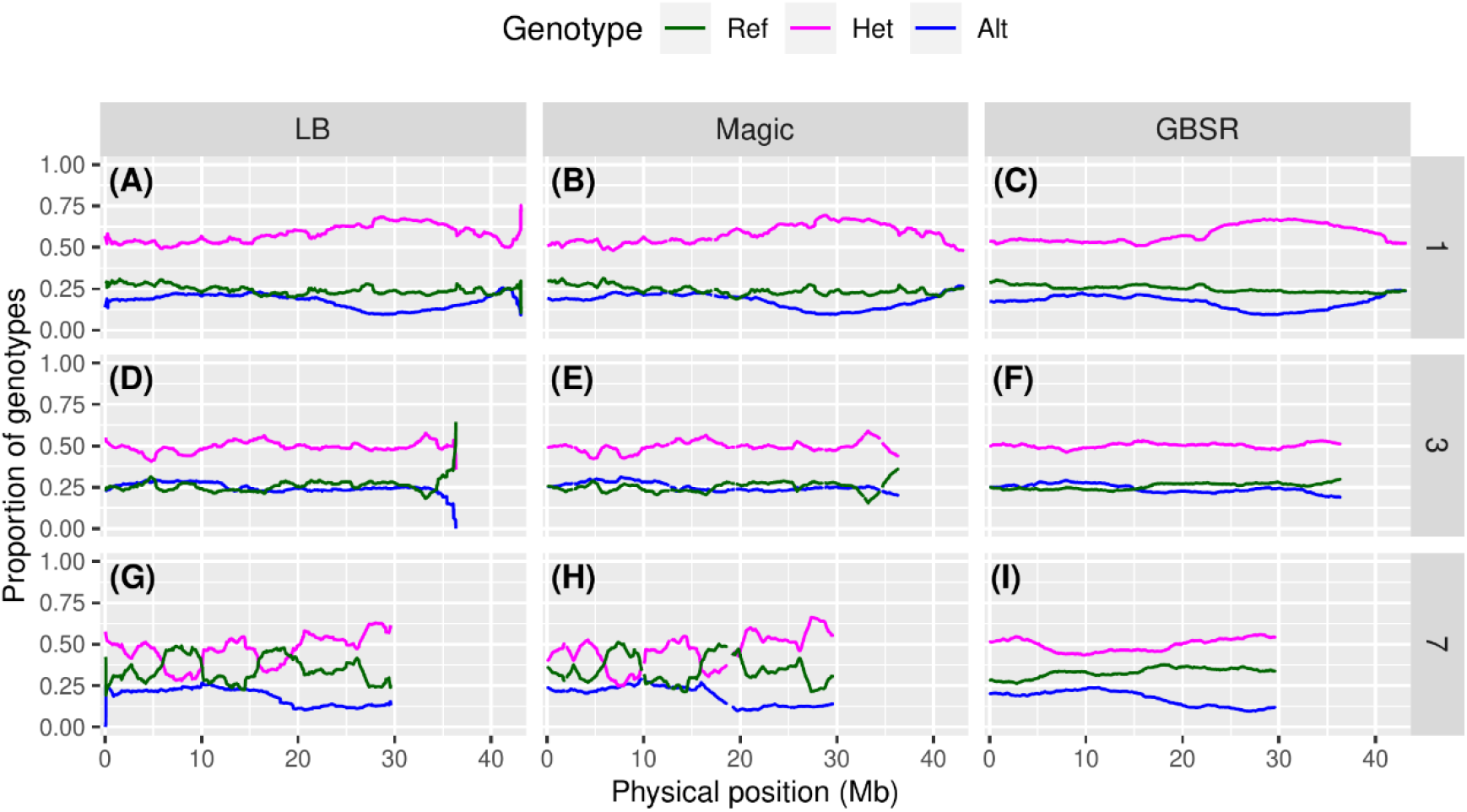
Genotype ratio of estimated offspring genotypes in the real data. Genotype ratios at markers along chromosome 1 (A-C), 3 (D-F), and 7 (G-I) in the real data are shown as line plots. “LB”, “Magic”, and “GBSR” indicate LB-Impute, magicImpute, and GBScleanR, respectively. Proportions of the reference homozygous, heterozygous, and alternative homozygous genotypes are represented by green, magenta, and blue lines, respectively.

### Comparing the running times

In addition to the comparisons of the algorithms in the aspects of accuracy and robustness, we also compared the running times using the simulation datasets. As shown in Table 2, GBScleanR completed all processes of genotype estimation faster than LB-Impute and magicImpute for the simulation datasets of biparental F_2_ populations including homoP2_F2 and hetP2_F2. In the case of the largest homoP2_F2 dataset that consists of 1000 samples, GBScleanR took only 15 seconds even with four cycles of IPO, while magicImpute and LB-Impute consumed 6.5 and 122.3 times as long. Although the magicImpute algorithm could only work properly for the hetP2_F2 datasets with detailed pedigree information given by a pedigree file, while the algorithm should accept simple specification of pedigree information given by a string like “Pairing, Sibling” indicating mating scheme of a given population (See *Input parameter settings* in the Materials and Methods section), we evaluated running times of magicImpute with or without pedigree files for hetP2_F2. Even for the hetP2_F2 datasets, which require a larger number of calculations than homoP2_F2, GBScleanR with IPO outperformed magicImpute (Table 2). Detailed pedigree information given as a pedigree file made magicImpute quite slower than that with simple pedigree information. On the other hand, a 2.5-times longer calculation at maximum was observed in GBScleanR with four cycles of IPO compared to magicImpute for the homoP8_RIL datasets (Table 2). Nevertheless, IPO was not required basically for RIL populations as shown in Figure 2, and GBScleanR without IPO could finish the calculations 35% faster than magicImpute even in the homoP8_RIL dataset with 1000 samples.

**Table 2.** Running time.

| Dataset | No. of samples | GBScleanR<sup>a</sup> | magicImpute<sup>b</sup> | LB-Impute |
| --- | --- | --- | --- | --- |
| homoP2_F2 | 10 | 2 (1) | 2 | 20 |
| homoP2_F2 | 100 | 3 (1) | 13 | 178 |
| homoP2_F2 | 1000 | 15 (5) | 97 | 1834 |
| hetP2_F2 | 10 | 4 (2) | 3 (10) | NA |
| hetP2_F2 | 100 | 18 (5) | 20 (140) | NA |
| hetP2_F2 | 1000 | 143 (41) | 165 (7447) | NA |
| homoP8_RIL | 10 | 44 (12) | 98 | NA |
| homoP8_RIL | 100 | 235 (62) | 336 | NA |
| homoP8_RIL | 1000 | 2087 (535) | 829 | NA |

Running times in seconds for each dataset are shown.
<sup>a</sup> The numbers in parenthesis indicate the running times of GBScleanR without the iterative parameter optimization.
<sup>b</sup> The numbers in parenthesis indicate the running times of magicImpute with pedigree files.

## Discussion

### Advantage of GBScleanR

The present study introduced an algorithm to provide robust error correction on genotype data with error prone markers that show biased probabilities for allele read acquisition. Any genotype data derived from GBS, or other RRS-based genotyping techniques, has a chance to contain such error prone markers even after trying to filter out erroneous markers. Particularly, a population derived from a cross between distant relatives, as in our case in which a cultivated rice *O. sativa* and a wild rice *O. longistaminata* were crossed, may potentially have a relatively high proportion of error prone markers. If a model uniformly assumes a 50% chance of obtaining a read for either allele for all markers at a heterozygous site, any HMM-based methods and sliding window methods can theoretically be deceived by the error prone markers.

Our simulation studies showed the advantage of using GBScleanR to precisely estimate the true genotype of noisy GBS data by incorporating the unevenness of allele reads into the model. Although all of the algorithms resulted in poor estimation accuracy for the simulation datasets with less than 1× of the offspring read depth, in which correct genotype call rates were smaller than 80%, especially in the hetP2_F2 datasets, GBScleanR with IPO basically outperformed the other algorithms in any situation and resulted in the most stable and accurate genotype estimation. When the offspring reads more than 3× could be provided for a population derived from inbred founders, including homoP2_F2 and homoP8_RIL, GBScleanR made nearly 95-99% accurate estimation (Figure 1 and Supplemental Data 1). On the other hand, biparental populations derived from outbred founders (hetP2_F2) seemed to require a larger number of reads for precise estimation. In addition, the number of samples in the given population exhibited a relatively large effect on the estimation of founder genotypes and the accuracy of founder genotype estimation affected on the accuracy of offspring genotype estimation (Figure 1 and 2). Our simulation study presented the difficulty of genotype estimation of a population derived from outbred founders compared with that derived from inbred founders. Interestingly, the increase of offspring reads could not improve genotype estimation accuracy for nosy GBS data by LB-Impute and magicImpute and rather decreased accuracy with the increase of miscall rates (Figure 1 and Supplemental Figure 2). Since GBScleanR without IPO also showed similar results with LB-Impute and magicImpute, those unexpected increases of miscalls were caused by biased genotype estimation on error prone markers that had uneven read acquisition ratio of reference and alternative alleles. The larger number of reads per maker would be expected to increase the reliability of genotype calling. However, in the case of genotype data with error prone markers, our study demonstrated that the algorithms without concern for allele read biases would mess up the genotype estimation.

The algorithm evaluation using the real data that potentially contains error prone markers also exhibited the superiority of GBSclenaR. The result clearly illustrated the negative impact of error prone markers having sever allele read biases and how the iterative estimation of error pattern by GBScleanR outperforms the existing algorithms (Figure 5 and Supplemental Figure 6). The estimated genotype data for the real data by LB-Impute resulted in the lowest missing rate and the highest recombination frequency compared with the other algorithms (Figure 4). Together with the relatively poor correct call rate in the simulation data (Figure 2), the LB-Impute algorithm tends to try filling up genotype calls even those are relatively less reliable and results in biased genotype estimation around error prone markers. Therefore, LB-Impute may deteriorate genotype data quality by enhancing wrong genotype calls using erroneous information of biased read counts if given genotype data contains error prone markers. Unlike LB-Impute, the magicImpute algorithm resulted in relatively small numbers of recombinations, while it scored the highest missing rates in all chromosomes (Figure 4). Although magicImpute seems to set missing to unreliable estimation, biased allele read ratios at error prone markers cannot be properly handled in the algorithm. As the result of biased genotype estimation around the error prone markers, magicImpute also resulted in highly fluctuating patterns of genotype ratio throughout almost all chromosomes, like LB-Impute did. On the other hand, GBScleanR could score the best concordance rates in all chromosomes (Figure 4). In addition, although the numbers of recombinations found in each chromosome did not fit the expected numbers according to the genetic map of rice estimated in a previous study^32^, the lowest number of the improbable double recombinations within short stretches was observed in the genotype data estimated by GBScleanR (Table 1). These results, together with the higher accuracy in the simulation datasets, indicates that GBScleanR is able to precisely incorporate the information of biased read counts at error prone markers and give robust and accurate estimation. The plots of the estimated genotype ratios also illustrate that GBScleanR can largely reduce the negative impacts of biased read counts and minimize artificial distortions even in the chromosomes containing a relatively large number of error prone markers (Figure 5 and Supplemental Figure 6).

The distortion level and the recombination frequency have large effects on downstream genetic analyses and also those values themselves are research targets. Any genetic and genomic study relies on the given genotype data and the accuracy of genotyping has a crucial effect on the quality of such research. The accuracy of genotyping can be evaluated by three criteria: correct calls, miscalls, and missing calls. Usually, miscalls have a more destructive effect on post-genotyping analyses than missing calls, because post-genotyping analyses regularly suppose all genotype calls to be true even if those include miscalls, and potentially generate misleading results. Furthermore, if the wrongly genotyped markers are clustered in a particular region, all statistical calculations for that region will be unreliable. Our study demonstrated the reliability of GBScleanR as the support tool for genetic studies that handle genotype data even with or without error prone markers.

### GBScleanR requires no strict filtering

In this study, we conducted a filtering based on a quantile of read counts for the real data to filter out genotype calls supported by overrepresented sequences that are possibly derived from paralogous and repetitive sequences. No one is able to exactly distinguish which reads were wrongly mapped. The error correction algorithms basically “believe” that the genotype calls supported by the higher number of reads were more probable, while fewer reads give the algorithms more space to guess the true genotype. The large number of reads with overrepresented mismapped reads does not increase the reliability of the genotype calls but rather causes misestimation of homozygous calls as heterozygous. Although GSBcleanR tries to detect mismapped reads, we recommend to filter out genotype calls supported by overrepresented sequences to reduce their negative impact on estimation. Nevertheless, GBScleanR accepts the error prone markers in a given dataset and recognizes the error pattern via IPO. Therefore, no strict filtering of markers based on allele frequency and missing rate is required, which is regularly performed for RRS-based genotyping data^33,34^. These filtering steps usually remove a large portion of the detected SNP markers^35–38^. In the case of our real data, filtering to remove possible error prone markers based on, for example, minor allele frequency > 5% and missing rate < 20%left only 162 out of 5035 SNPs for the 12 chromosomes in the real data. Markers that would be filtered out may possibly contain information about the true genotype at the markers even if the data are partial and skewed. GBScleanR is able to integrate those less informative markers into the error correction process by tweaking the HMM. Since there is no clear guideline for filtering strength, which should differ depending on the dataset used, filtering criteria are left in the hands of researchers. GBScleanR allows users to skip such an arbitrary step and improves the cost efficiency of the RRS-based genotyping system by reducing the number of unused reads obtained via costly NGS.

### Limitations and perspectives of GBScleanR

The current implementation of GBScleanR treats all chromosomes as diploid autosomes. Although sex chromosomes can be processed by our algorithm, the calculation of transition probabilities between autosomes and sex chromosomes would be expected to differ, in which any recombination does not occur in either male or female individuals. The algorithm therefore overestimates the number of recombinations in the sex chromosomes, while Supplemental Figure 8B shows that an overrated recombination frequency does not produce any severe effects on genotype estimation. A previously published paper introduced a model that can be used to infer the number of recombinations in the sex chromosomes of experimental populations^39^. We plan to further implement this model into GBScleanR in a future update.

In addition, there are obvious demands for applying GBScleanR to polyploid organisms. Several dosage estimation algorithms have already been implemented some tools^40–42^. Nevertheless, all of these infer the dosage at each marker independently and do not incorporate information at adjacent markers, as performed in an HMM. The dosage estimation using an HMM may increase the robustness against error prone markers, while increasing the complexity of the algorithm at the same time. The codes of GBScleanR have been basically organized to allow us to expand its functions for polyploid organisms. Thus, GBScleanR will support polyploids in the future.

The recent remarkable development in third-generation sequencing, named Sequel by Pacific Bioscience and MinION by Oxford Nanopore Technologies, enables us to obtain high quality genome sequences at reasonable cost^43,44^. Reference genome information for any organism is expected to become available soon. Thus, RRS-based genotyping systems that are underpinned by a flexible robust error correction tool such as GBScleanR will become practical platforms for use in breeding programs that require a prompt and efficient scheme under limited periods and budgets. Distant relatives including the ancestral wild species of domesticated species are important genetic resources, particularly in plant breeding, that can improve abiotic and biotic stress resistance^45–47^. For such genetic studies and breeding projects using wild species, GBScleanR could be the first choice as a fundamental tool for genotype identification coupled with an RRS-based genotyping system.

## Methods

### Overview of the GBScleanR package

GBScleanR is an R package and consists of a main tool that conducts error correction and utility functions that handle genotype data in the variant call format (VCF). Supplemental Figure 7 presents the data analysis work-flow and the related functions that are implemented in GBScleanR. The algorithm is designed to estimate the true genotype calls along chromosomes from the given allele read counts in a VCF file that is generated by SNP callers such as GATK and TASSEL-GBS^48,49^. The current implementation supports genotypic data for mapping populations that are derived from two or more diploid founders followed by selfings, sibling crosses, or random crosses, for example F_2_ and 8-way RILs. Our method assumes that markers are biallelic and ordered along the chromosomes, and the reads are mapped onto a reference genome sequence. To access the large amounts of genotype data required, an input VCF file is first converted into a genomic data structure (GDS) file^50^. This conversion automatically filter out non-biallelic markers.

GBScleanR provides functions for data visualization, filtering, and loading/writing a VCF file. Furthermore, the data structure of the GDS file created via this package is compatible with that used in the SeqArray package and can be further converted to a derivative GDS format used in the SNPRelate, GWASTools and GENESIS packages, which are designed to handle large amounts of variant data and conduct regression analysis^50,52–54^. Therefore, GBScleanR can be built in a pipeline for an genetic study that allows us smooth access to the given large genotype data for visualization, filtering, error correction, and regression analysis without the requirement for extra data format conversion steps.

### Modeling for error correction

The error correction algorithm of GBScleanR is based on the HMM and treats the observed allele read counts for each SNP marker along a chromosome as outputs from a sequence of latent true genotypes. Our model supposes that a population of *N*° ≥ 1 sampled offspring is originally derived from crosses between *N*^f^ ≥ 2 founder individuals. The founders can be inbred lines with homozygotes at all markers or outbred lines in which markers would be heterozygous. Only one chromosome is considered for modeling due to the independence of chromosomes. Let 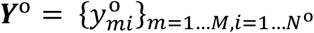 denote the observed allele read counts at marker *m* in offspring *i*. The element 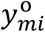 consists of two values, *y*^ref^ and *y*^alt^, which represent the reference read count and the alternative read count, respectively. Similarly, the observed allele read counts at marker *m* in founder *j* are represented by 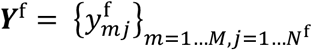. On the other hand, the matrices for hidden true offspring genotypes and hidden true founder genotypes are represented by 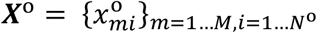 and 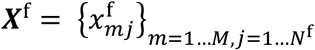, respectively. The element 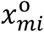, takes a value of 0, 1, or 2 to indicate the reference homozygote, heterozygote, and alternative homozygote genotype without phasing information. Unlike the offspring genotype matrix, the founder genotype 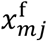 stores the phased genotype as 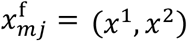 where *x*^1^ and *x*^2^ indicate alleles at marker *m* on one of the diploid chromosomes and another in founder *j*, respectively. The reference allele is represented by 0, while 1 denotes the alternative.

Considering the linkage between the markers and the independence of the founder genotypes, we can assume that the sequence of the genotypes in the *i*^th^ sample, which is denoted as 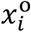, does not follow a Markov process, while the sequence of the descendent haplotypes 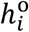 along a pair of chromosomes that is derived from *N*^f^ founders does. To enable our estimation problem to be solved in the HMM framework, 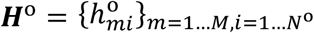 is introduced to represent the matrix of the phased descendent haplotypes. The element 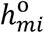 denotes a pair of descendent haplotypes (*h*^1^, *h*^2^) at marker *m* in offspring *i*. Therefore, *h*^1^ and *h*^2^ each takes one of the natural numbers that are ≤ *N*^f^ if all founders are inbred or ≤2*N*^f^ if outbred, to indicate the origins of the descendent haplotypes.

The algorithm estimates ***H***^o^ and ***X***^o^ based on ***Y***^o^ and ***Y***^f^ by maximizing the joint probability *P*(*H*^o^, ***X***^f^, ***Y***^o^, ***Y***^f^). The parameters used in the model are not shown here. The joint probability can be transformed as seen below.

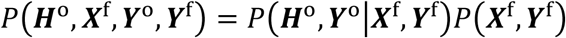

Each probability on the right hand of the equation can be farther transformed.

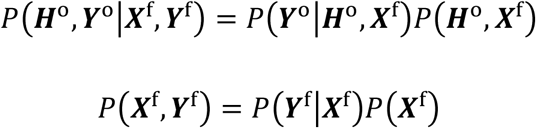

We assume the independence of the offspring and then rewrite the first equation as

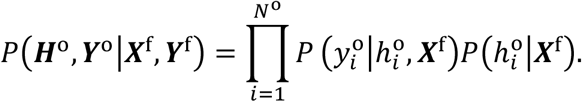

We can now treat 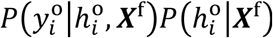 as the HMM for *i^th^* offspring’s haplotypes and the read counts, given the founder genotype ***X***^f^. On the other hand, *P*(***Y***^f^|***X***^f^)*P*(***X***^f^) is the product of the probability of observing ***Y***^f^ under the condition ***X***^f^ and the probability of obtaining ***X***^f^. The founder genotypes at each marker can be considered independent of each other and do not follow a Markov process. Therefore, our algorithm tries to maximize the following joint probability:

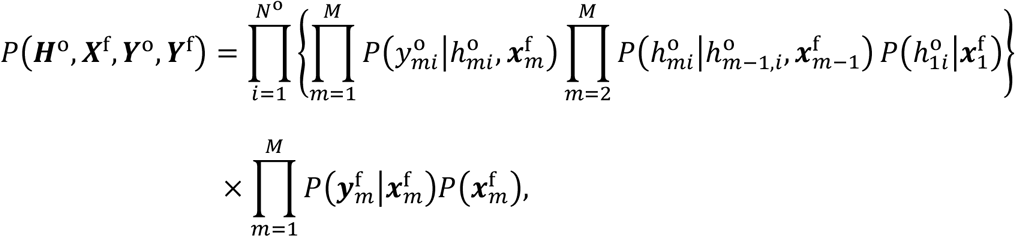

where 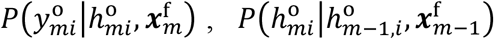 and 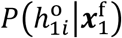 correspond to the emission probability, the transition probability, and the initial probability of the HMM, respectively, and 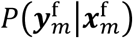 is the probability of observing read counts for the founders at marker *m* when the combination of the true genotypes of the founders is 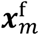. We assume that all SNP markers are biallelic and that 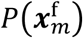 follows a discrete uniform probability for all possible combinations of the founder genotypes, while omitting cases in which all founders have homozygotes on the same allele, e.g., when all founders have reference homozygote genotypes.

### Emission probability

The descendent haplotype 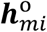 itself does not provide any information about the alleles that are obtained by the descendants of the founders. We can only calculate the probabilities of obtaining 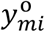 from 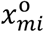, but not from 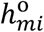 directly. Therefore, to obtain the emission probability 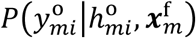, three additional probabilities are introduced; 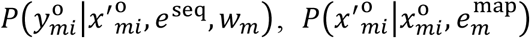, and 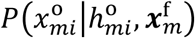, where *e*^seq^ and 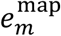 represent the sequencing error rate and the mapping error rate of the observed reads, respectively. The allele read bias *w_m_* is the key parameter of the HMM in GBScleanR that accounts for systematic bias toward either of the alleles in the reads. We assume that each marker has its own mapping error rate and allele read bias, while the sequencing error rate is set to be uniform for all markers. 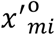 indicates the observable genotype that results from accounting for the mismapping of reads with a mapping error rate of 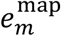. Combining these parameters allows the emission probability to be replaced with 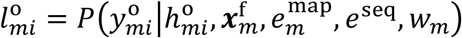 and it holds that

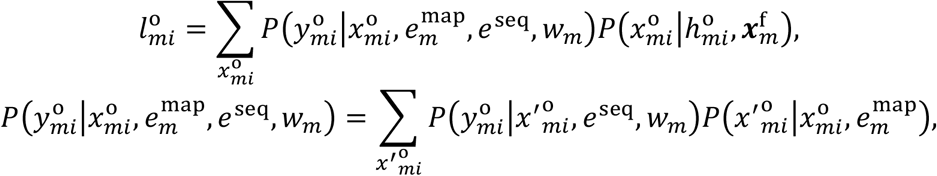

where 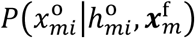 is given a value of 1 for the possible genotype under the constraint of the descendent haplotypes and founder genotypes in sample *i* and 0 for the other genotypes. For example, if sample 1 has 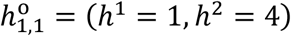 at marker 1, indicating that the haplotypes descended from the first chromosome of founder 1 and the second chromosome of founder 2 and if the founder genotypes are 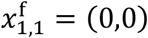 and 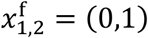, the genotype can only be heterozygotic 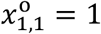.

We assume that the observed read counts 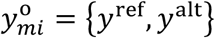 follow binomial distribution

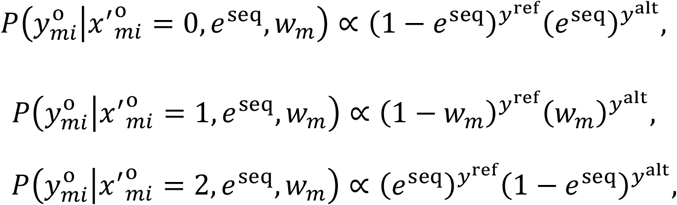

given the observable genotype 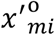. Where no bias is assumed in the allele reads, *w_m_* = 0.5, is used as in the previously published models^24,26,27^. The normalization constant for 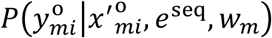 is not shown in the equation above. In addition to sequencing errors and allele observation bias, read mismapping is also a well-known error form in NGS based genotyping. Mismapping leads to the false observation of reads and leads to false heterozygote detection when the true genotype is homozygous. This type of error is accounted for by summing over 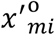 in the joint probability 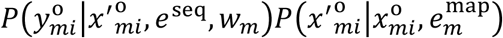. We assume that 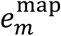 can take two values (*e*^ref^, *e*^alt^), indicating the probability of observing mismapped alternative reads when the true genotype is the same as the reference homozygote and mismapped reference reads when the true genotype is an alternative homozygote. Thus, the probability 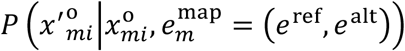 takes the values listed in Table 3.

**Table 3.**
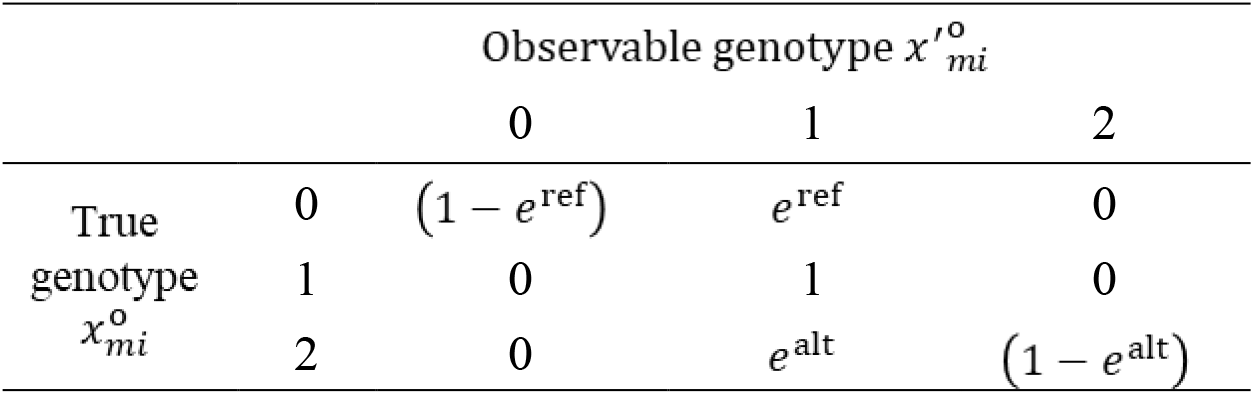
Probabilities associated with mismapping.

Similar to the emission probability for offspring, the emission probability for founders can be denoted as 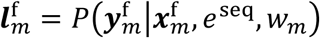 that represents the probability of observing the given read counts in the founders 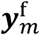 given a combination of the true founder genotypes 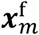. It is obtained by

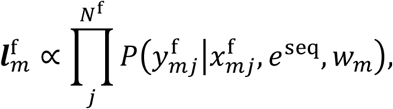

where the mismapping of reads in the founders is not considered. The normalization constant for 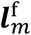 is not shown. To omit the less probable founder genotypes, the probability 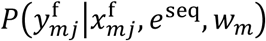 is set at 0 if the result is less than 0.01.

### Transition probability

The transition probability 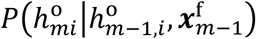 is used to describe the probability that a crossover occurs between adjacent markers. The sequences of the descendent haplotypes are modelled using a continuous-time Markov chain, as described previously^55,56^. The probability that the haplotype state transitions from 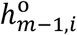 to 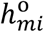 between the markers can be obtained using

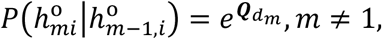

where **Q** denotes the transition-rate matrix that defines the rates at which a state transition occurs per unit of time.

The parameter **Q** is calculated based on the idea of junction densities following the modelling framework described in Zheng *et al*., 2014 and Zheng *et al*, 2015^56^. The junction densities define the expected numbers of junction types per Morgan. If *j*(*abcd*) represent a junction density of the type (*abcd*) where *ab* indicates a pair of descendent haplotypes at a marker and *cd* denotes that of the next marker located at the other side of the junction. Each of *a, b, c, d* can take one of the integers indicating the origin of the haplotype. j(1112) indicates one junction from haplotype 1 to 2 on one chromosome, with the same haplotype maintained on the other chromosome. In the case of a biparental population derived from inbred founders, possible junction types are j(1111), j(1212), j(2222), j(1112), j(1211), j(1221), j(1122), and the junction types in which the order of numbers are swapped, e.g j(2121). As genetic markers are considered sufficiently dense in RRS-based genotype data, only one junction between adjacent markers is expected at most. Therefore, the rate in the transition-rate matrix **Q** was set to 0 for the junction types with more than one junction e.g., j(1221) and j(2112). Although j(1122) also has two junctions, these could be observed in a later generation because the chromosomes are both derived from one identical chromosome that had a junction between haplotypes 1 and 2 in a preceding generation. The possible junctions would be selected based on pedigree information. There is no chance that a third haplotype, e.g., j(1213), can be obtained in a biparental population, but it is possible in a population that is derived from four or more founders, such as 8-way RILs.

The matrix exponential is obtained by Higham’s algorithm, which is implemented in the “expm” package of R (http://cran.r-project.org/web/packages/expm/index.html). To obtain transition rates that are proportional to the distances between adjacent markers, **Q** is multiplied by the genetic distance between markers *m* and *m-1* in Morgans *d_m_*. While our algorithm requires genetic distances between markers, genotype data obtained via NGS generally provides only information about the physical distances. Therefore, *d_m_* is calculated based on the physical distance and the expected genetic distance:

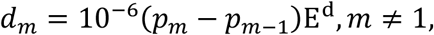

where *p_m_* represents the physical position of marker *m* and E^d^ is the expected genetic distance per mega base pairs that should be specified as input parameter.

For m = 1, the initial probability 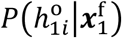 is required and specified by the stationary distribution of the Markov process that can be obtained by normalizing the first eigenvector of the transposed transition probability for a marker interval.

### Estimation of the best haplotype and genotype sequences

Together with the definitions described in the sections above, the joint probability that we need to maximize can be rewritten as shown below.

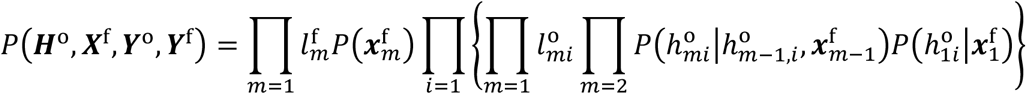

The Viterbi algorithm is used to maximize the joint probability and estimate the most probable haplotype sequences^57^. Considering that the founder genotype itself does not follow the Markov process while the offspring haplotypes depend on the founder genotypes, the most probable founder genotypes at marker *m* can be estimated based on the scores (probabilities) and the states of the Viterbi trellises that were calculated for the offspring genotype from marker 1 to *m*. Therefore, our algorithm recursively calculates the Viterbi scores for offspring 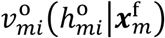 and founders 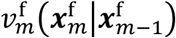, and records the Viterbi paths for the sequences of offspring haplotypes 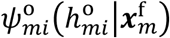 and founder genotypes 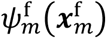, respectively. The recursive calculation via an integrated Viterbi algorithm proceeds as follows.

Initialization at m = 1:

The initialization starts with simply calculating the probability to have 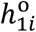 when the combination of the founder genotypes 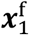 was given.

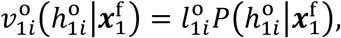

where 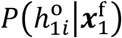 is the initial probability that the *i^th^* offspring has the combination of haplotypes 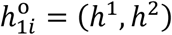 at marker m.

Recursion at 2 ≤ m ≤ M:

The recursive calculation can be separated into several steps. As the first step, the algorithm calculates the probabilities of state transitions from marker m-1 to m as shown below.

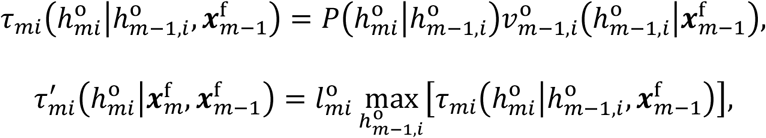

where 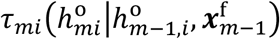 is a matrix each element of that indicates the probability to observe the transition from 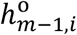 to 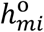 when 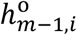 and 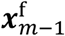 are given, while 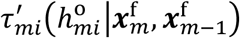 indicates the probabilities to observe 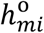 when the most probable transition from 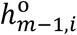 to 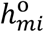 occurred with given 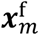 and 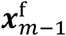.

Then, calculate the Viterbi scores and the Viterbi paths for founders as

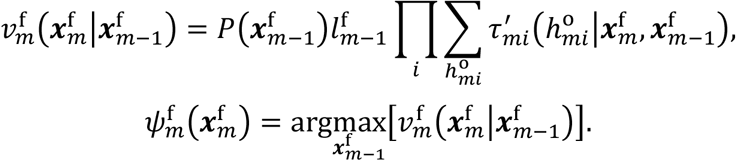

Using 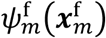 that stores the most probable combination of founder genotypes at marker m-1 when 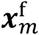 is given, we can get the Viterbi scores and the Viterbi paths for offspring as following.

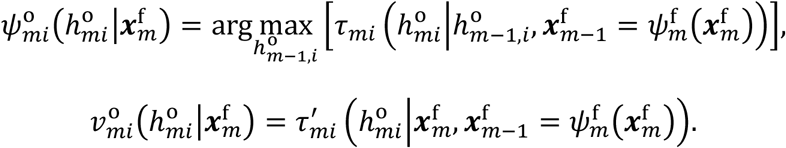

Termination at m = M:

Finally, we obtain the probability to observe 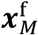 and the most probable combination of founder genotypes at marker M via the following calculation.

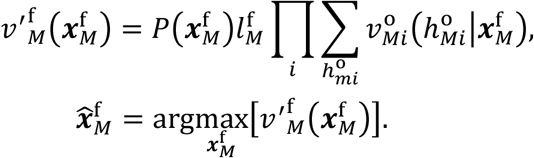

Once 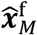 was determined, we can also get the most probable offspring haplotypes at marker M:

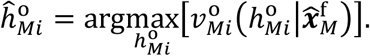

Backtracking from M-1 to 1:

To retrieve the most probable sequences of founder genotypes and offspring haplotypes, we trace back the Viterbi paths as shown below.

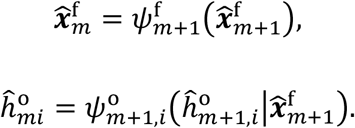

If multiple arguments have the same maximum score, one of them is randomly selected. To increase the accuracy, our algorithm executes two rounds of estimation: one from marker 1 to M and one in the reverse direction. The reverse direction round starts from the last marker M and uses the estimated founder genotypes 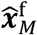 from the forward round as the only possible founder genotypes at marker M. The results from both rounds are then merged by combining the first half of the estimated genotypes 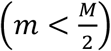 from the reversed direction round with the last half from the forward direction round.

The estimated genotype 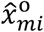 at each marker in each sample can now be obtained based on 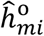 and 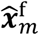. To evaluate the reliability of the estimated genotypes, the marginal probability is also calculated:

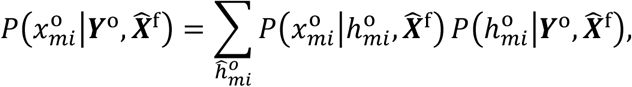

where 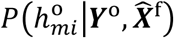 can be obtained using the forward-backward algorithm^57^. The estimated genotype 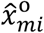 is set as missing if 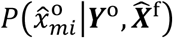 is smaller than a user specified threshold Pcall, with 0.9 as the default.

### Estimation of allele read bias and mismapping rate

GBScleanR requires four parameters E^d^, *e*^seq^, 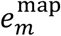, and w_m_. The expected genetic distance per mega base pairs E^d^ and sequencing error rate *e*^seq^ take a single positive number that should arbitrarily be specified as input parameters, while the mismap rates 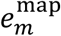 and the allele read biases *w_m_* can be estimated via the iterative parameter optimization (IPO). To obtain precise estimations of the parameters, the algorithm iteratively performs genotype estimation via the Viterbi algorithm, as described in the previous section, and parameter estimation via the procedure described here. For the first cycle of genotype estimation, 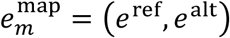, and *w_m_* are initialized to (0,0) and 0.5 for all markers, respectively. Thus, the first iteration does not consider allele read biases and mismapped reads, although *e*^seq^ can partially account for mismapped reads. With the estimated genotypes from the first cycle, we can obtain the numbers of reads for each estimated genotype. Let 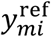 and 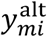 be the number of reference reads and alternative reads at marker *m* in sample *i* including both offspring and founders. The mismapping rate at marker *m* can be obtained using:

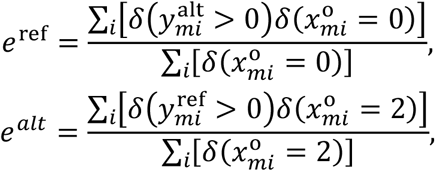

where *δ* is an indicator function that is equal to 1 if the argument is true and 0 if it is false.

Similarly, the allele read bias at marker *m* is estimated by the following:

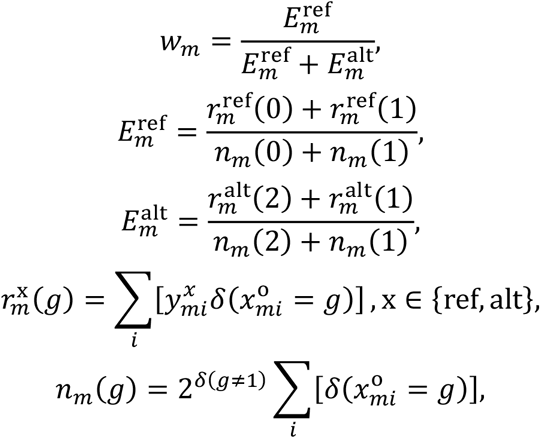

where *g* is 0, 1, or 2 and indicates the genotype; 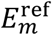 represents the expected number of reference allele reads observed if one reference allele is on the chromosome, and 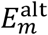 represents the expected number of alternative allele reads if one alternative allele is on the chromosome. This calculation relies on the assumption that the reads at each marker in both heterozygous samples and homozygous samples can be biased if samples are prepared and sequenced together. To confirm if the given data meets this assumption, our algorithm first calculates ***w*** = {*w_m_*}_*m*=1,…*M*_ with *g* = 1 and *g* = {0,2} separately, and then evaluates the correlation between ***w*** obtained in both calculations. If the correlation coefficient is greater than or equal to 0.7, the equation above is employed; otherwise, ***w*** used with g = 1, although all of the datasets used in this paper meet the assumption of bias uniformity, even the real dataset.

### Preparation of real data

The genotype data of an F_2_ population that was derived from a cross between *O. sativa* and *O. longistaminata* were used in this study. The data potentially include a large number of error prone markers with mismapping and allele read biases, as described in our previous paper^58^. In brief, the F2 population was produced via the self-pollination of F_1_ plants that were derived from a cross between *O. sativa* ssp. *japonica* cv. Nipponbare and *O. longistaminata* acc. IRGC110404. GBS was performed with a *Kpn*I-*Msp*I restriction enzyme pair using MiSeq with 75 bp paired-end sequencing. Sequencing runs with nine 96-multiplex libraries generated an average 134,447 ± 50,788 reads per sample. We obtained a total of 2,539,459 and 3,481,218 reads for *O. sativa* and *O. longistaminata* from the independent runs, respectively, which should be sufficient for fairly accurate determination of the founder genotypes. The obtained reads were then processed via the TASSEL-GBS pipeline v2 by following the instructions in the manual with the default parameters (https://bitbucket.org/tasseladmin/tassel-5-source/wiki/Tassel5GBSv2Pipeline)^49^. The obtained SNP markers were then filtered to retain only those markers that were homozygous in each founder but biallelic between founders. Only the first SNP was selected if multiple SNPs were present within a 75 bp stretch. Considering the distant cross that was used to obtain the F_2_ population, our genotype data might include mismap-prone markers due to the repetitive sequences in the two founder genomes. The mismapping patterns can differ between samples which have different descendent genome compositions. Therefore, to filter out the erroneous genotype calls that were caused by mismapping reads from repetitive sequences, both the reference and the alternative allele read counts were set to 0 for genotype calls where either the reference or the alternative allele reads were higher than the 90th percentile for each genotype call in each sample, respectively. The resulting genotype data had 5035 SNP markers for 814 F2 individuals with an average read depth of 0.85×.

### Generation of simulation data

Genotype datasets were simulated for three scenarios: a 2-way F_2_ population from inbred founders (homoP2_F2), a 2-way F_2_ population from outbred founders (hetP2_F2), and 8-way RILs from inbred founders (homoP8_RIL). For each scenario, we simulated populations that have 10, 100, and 1000 individuals, respectively. To generate the descendent haplotype and genotype data for each dataset in each scenario, the breeding pedigree was first simulated via the following. The homoP2_F2 scenario assumes that the two inbred founders have homozygotes at all markers and are biallelic between founders, the founders are crossed and then self-pollinate, as usually conducted in the genetic analysis of self-fertilizing plants. The hetP2_F2 scenario assumes two outbred founders with independently random genotypes at any marker with the minor allele observed in at least one of the four possible haplotypes. These outbred founders are subjected to mating to produce 1000 progenies and then 100 sibling combinations are used to produce 10 progenies in each. The last generation of hetP2_F2 has 1000 offspring in total. This scenario assumes the case of genetic analyses for non-self-fertilizing organisms. The pedigree of homoP8_RIL consists of three pairing generations and five self-fertilized generations. In the pairing stage, four pairs are selected from the eight founders with no duplication, and one descendant is created from each pair. The four descendants are then further paired to produce 100 progenies from each of the two pairs. At this step, we have two sibling families with 100 individuals in each. At the last pairing, 50 pairs are produced by selecting one individual per sibling family to produce pairs from each of which 50 progenies are generated. The resultant 2500 individuals are then self-fertilized five times so that one offspring is produced per individual in order to maintain the population size. This kind of pedigree is usually found in populations that have been subjected to genetic survey and breeding and known as Multiparent Advanced Generation Inter-Cross (MAGIC) population. A given number of offspring is sampled from the last generation to prepare each dataset.

Descendent haplotype data from each population was generated by introducing crossovers into the gametes that have descended from the parent(s) of each progeny at each mating step. Only one pair of 50 Mb chromosomes as diploid genome was simulated for all datasets and the expected genetic distance per mega base pairs was set to 0.04. Markers were located randomly throughout the chromosome. The occurrence of crossovers between adjacent markers in a gamete follows Poisson distribution with a mean value that is equal to the genetic distance of the given pair of adjacent markers. The genetic distances between the markers were calculated by multiplying the physical distances by 0.04×10^−6^. If an odd number of crossovers was assigned to a marker pair, gamete switching descendent haplotypes were generated, while the assigned crossover was ignored if the donor parent of the gamete showed IBD around the markers. The offspring genotypes were obtained by replacing the descendent haplotype at each marker with the genotype that was assigned to the founder haplotype.

Based on the simulated genotype data, read counts were assigned to each marker of each individual. The expected offspring read depths were set at 0.1×, 0.25×, 0.5×, 0.75×, 1×, 2×, 3×, 10×, and 20×, respectively. The founder read depth was set at 5× for all datasets, because usually founders are sequenced relatively deep even in the low coverage GBS. For each of the three scenarios, the read count data was simulated by mimicking the allele read biases observed in the real data. Generally, the total number of reads in read count data that is obtained via NGS is competitively allocated to markers of sequenced samples with biases that depend on the proportion of sequencing targets in a sequencing library. To mimic the competitiveness of reads between markers of samples, reads were allocated following the joint probability for all markers and samples as explained below.

The total number of reads per marker 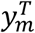 is assumed to follow exponential distribution with the rate parameter λ, Exp(*λ*). The probability of a read at marker *m* can be obtained by the following:

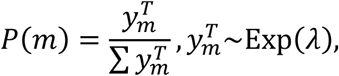

where λ is set at 0.25. The read obtained at marker *m* can be either reference or alternative with probabilities of p^ref^ and (1 – *p*^ref^), respectively, where *p*^ref^ represents the probability that the read is a reference allele. The probability *p*^ref^ takes different values depending on the genotype at marker *m* in sample *i, x_mi_*. When the genotype is the homozygous reference, *p*^ref^ = (1 – *e*^seq^), where *e*^seq^ represents a sequencing error. Similarly, *p*^ref^ = 0.5 and *p*^ref^ = *e*^seq^ for heterozygous and homozygous alternatives. To incorporate allele read bias, the genotype-dependent read observation probability *P*(*z*|*w_m_, x_mi_*), *z* ∈ {ref, alt} was introduced, which takes the values shown in Table 4. The joint probability of having a read for allele *z* at marker *m* in sample *i* can then be calculated:

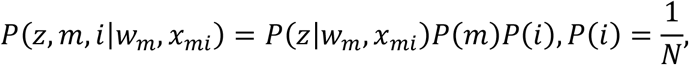

where *N* is the number of samples. Mismapping error is introduced by replacing the genotypes with the probability 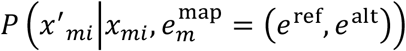, taking the values listed in Table 3. The number of total reads is calculated by multiplying the number of samples by the expected read depth and the number of markers. We then allocate those reads based on the joint probability *P*(*z,m, i*|*w_m_,x*′_*mi*_), which is not conditional on the true genotype *x_mi_*. The read count data for the founders and the offspring are simulated separately with given expected read depths for each dataset.

**Table 4.** Genotype-dependent read observation probabilities.

| | | Observable allele read $z$ | |
| --- | --- | --- | --- |
|  |  | Ref | Alt |
| Simulated<br>genotype<br>$x_{mi}$ | 0 | $(1 - e^{\text{seq}})w_m$ | $e^{\text{seq}}w_m$ |
| | 1 | $0.5w_m$ | $0.5(1 - w_m)$ |
| | 2 | $e^{\text{seq}}(1 - w_m)$ | $(1 - e^{\text{seq}})(1 - w_m)$ |

The simulation datasets were generated by mimicking only the real data of chromosome 7, which showed relatively severe bias (Supplemental Figure 1). Each dataset was therefore simulated to consist of 620 markers on one chromosome with physical lengths of 50 Mb and genetic lengths of 2 Morgans. *P*(*m*) was also obtained from the read data by calculating the total number of reads per marker 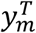 by summing up the read counts for each marker. The values *w_m_* were obtained by calculating 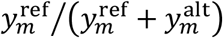, where 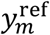 and 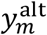 denote the total numbers of reference and alternative reads per marker observed in the real data, respectively. The mismapping rate was randomly assigned to markers by sampling the probabilities from an exponential distribution with the mean at 0.05, since the mismapping pattern in the real data is unknown. The sequencing error rate *e*^seq^ was set to 0.0025. Since typical genotyping pipeline filter out markers at which no read was observed in either of founders, we allocated at least one read to all markers of founders and then allocated the remaining reads by following the join probability (“nonzero” datasets). In addition to them, we also simulated datasets in which all reads were allocated by following the join probability and, as a result, founders have no read at some markers and missing genotypes (“allowzero” datasets).

### Input parameter settings

GBScleanR was tested and compared with LB-impute and magicImpute^26,27^. GBScleanR includes some arguments that can be used to tweak this tool: the expected genetic distance per mega base pairs E^d^, sequencing error rate *e*^seq^, threshold for the probability of calling estimated genotype P_call_, and the number of iterations required for genotype estimation. To decide the default values for these arguments, we tested the algorithm using the simulation dataset with 100 individuals and a 3× offspring read depth in the “nonzero” dataset of the homoP2_F2 scenario under various settings. As shown in Supplemental Figure S7, the tests indicated that P_call_ have relatively large impacts on the genotype estimation accuracy, while the differences associated with the number of iterations (if more than one), E^d^, and *e*^seq^ were negligible. The values of E^d^ = 0.04 and *e*^seq^ = 0.0025 that were used in the data simulation seem to be good choices when set for genotype estimation. Increasing P_call_ from 0.8 to 0.95 improved the correct call rate in an almost proportional manner, while the miscall rate worsened at the same time. Therefore, we set P_call_ at 0.9 for the algorithm evaluation as used in the default setting for magicImpute. In the case of iteration number, we found that a larger number leads to better correct call and miscall rates, which reached a plateau at four or more iterations. Thus, we decided to set the number of iterations to four. To evaluate GBScleanR, we tested the algorithm with and without the iterative parameters optimization (IPO) *w_m_* and 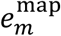.

For fair comparison among the three tools, similar input parameters were used if LB-Impute and magicImpute have corresponding parameters to those given in GBScleanR. In the case of LB-impute, the “impute” option was set to the “-method” argument and the “-offspringimpute” flag was used to run it in the offspring impute mode. A vector of strings stating the names of the founder samples was specified as “-parents”. “-recombdist” and “-readerr” were set at 12500000 and 0.0025, which correspond to E_d_ and *e*^seq^. While the parameter related to read mismapping is estimated by IPO and not tunable in GBScleanR, LB-Impute requires to set the parameter “-genotypeerr” that is related to *e*^map^ of GBScleanR. We set 0.005 to “-genotypeerr” as magicImpute also requires similar parameters and has 0.005 as default. The arguments in magicImpute include the mandatory arguments “model” and “popdesign” in addition to the genotype data in the “magicsnp” format. The simulated data and real data were converted to the magicsnp format. Marker positions in centimorgan, which are required for magicImpute, were calculated by multiplying the physical positions of the markers by the expected genetic distance per mega base of 0.04, which was used to generate the simulation data. The “jointModel” option was set to the “model” argument for all datasets. The “popdesign” argument that affects on the state transition-rates was specified based on the pedigree in each scenario as follows. The “popdesign” was set to (“Pairing”, “Selfing”) for the scenario homoP2_F2, while (“Pairing”, “Pairing”, “Pairing”, “Selfing”, “Selfing”, “Selfing”, “Selfing”, “Selfing”) was specified for homoP8_RIL. Although the popdesign for hetP2_F2 was supposed to be (“Pairing”, “Sibling”) based on the pedigree and the instructions in the manual, magicImpute did not accept it and returned an error message saying “invalid popdesign”. Therefore, pedigree information files were created to indicate population design for all datasets of the hetP2_F2 scenario and provided to execute magicImpute as “popdesign”. As a note, we found that the execution of magicImpute with a pedigree information file takes longer calculation than that with simple “popdesign” specification by a string vector. Additionally, magicImpute has “minPhredQualScore”, “imputingThreshold”, and “detectingThreshold” arguments. The former corresponds to *e*^seq^ and was set at 26.0206 that is equal to *e*^seq^ = 0.0025. The latter two specify the parameter similar with Pcall of GBScleanR and both of them was set at 0.9. Both of “founderAllelicError” and “offspringAllelicError” that corresponds to *e*^map^ of GBScleanR and “-genotypeerr” of LB-Impute were set to 0.005. The estimated genotype data generated by magicImpute includes incomplete genotype estimations represented by 1N, N1, 2N, and N2, where N points to an unknown allele and the numbers 1 and 2 indicate reference and alternative alleles, respectively. Although these incomplete estimations provide partial information about the genotype at the markers, we treat them as completely missing because downsream genetic analyses generally cannot handle such incomplete information.

### Evaluation of algorithms

The genotype estimation accuracy of each algorithm for the simulated datasets was evaluated by calculating the proportions of correctly estimated genotypes (correct call rate), wrongly estimated genotypes (miscall rate), and missing genotype calls (missing call rate) for each offspring in the datasets. The means and standard deviations of these values were then obtained for each dataset processed by each algorithm. The resultant scores are shown in Supplemental Data 1. In the case of the real data, we first selected relatively reliable genotype calls supported by more than 6 reads at markers that have less than 20% of missing rate and more than 40% of minor allele frequency. Those selected genotype calls were then set to missing by removing all read counts observed for them. This masked real data was subjected to genotype estimation using GBScleanR, LB-Impute, and magicImpute. The resultant estimated genotype data was evaluated based on the concordance rate that is the proportion of estimated genotype calls matching with the raw genotype calls at the masked genotype calls. Furthermore, we also calculated the averages of missing rate, the number of recombinations, and segregation distortion level in each chromosome. The segregation distortion level at each marker was obtained by dividing the chi-square value, which was calculated based on the assumption that the genotype ratio to be 0.25, 0.5, and 0.25 for three possible genotypes, by the number of non-missing genotype calls. The segregation distortion level of each chromosome was then obtained by summing up the segregation distortion levels of all markers in each chromosome. To identify recombination breakpoints of genome segments in the estimated genotype data, we searched the markers that show genotypes switched between adjacent non-missing markers. The middle points of those markers were considered as recombination break points.

## Data availability

The stable version of GBScleanR is available on Bioconductor (https://bioconductor.org/packages/GBScleanR/). The latest developmental version is available on GitHub (https://github.com/tomoyukif/GBScleanR). This package and its source codes are distributed under the license GNU General Public License version 3. The vcf file of the real data and R scripts for the data analyses performed in this study are also available on GitHub (https://github.com/tomoyukif/GBSR_SupFiles). Further description of the sequencing reads for the real data can be found in Furuta *et al*., 2017^58^.

## Acknowledgements

This work was supported by the CREST program of the Japan Science and Technology Agency [JPMJCR13B1 to M.A.]; the SATREPS program in collaboration of the Japan Science and Technology Agency and the Japan International Cooperation Agency [JPMJSA1706 to T.F. and M.A.]; KAKENHI of the Ministry of Education, Culture, Sports, Science and Technology of Japan [JP20H05912 to M.A.]; and KAKENHI of Japan Society for the Promotion of Science [JP20K15503 to T.F. and JP20H02958 to T.Y.].

## Contributions

T.F. conceived and designed the study. T.F. also developed the R package, conducted bioinformatics analysis, and drafted the manuscript. M.A. and T.Y. supervised the study. All authors read and approved the final version of the manuscript.

