## Supplementary material for "GBScleanR: Robust genotyping error correction using hidden Markov model with error pattern recognition": SupplementalMaterials.docx

*To whom correspondence should be addressed.

Supplemental Figures S1-S7

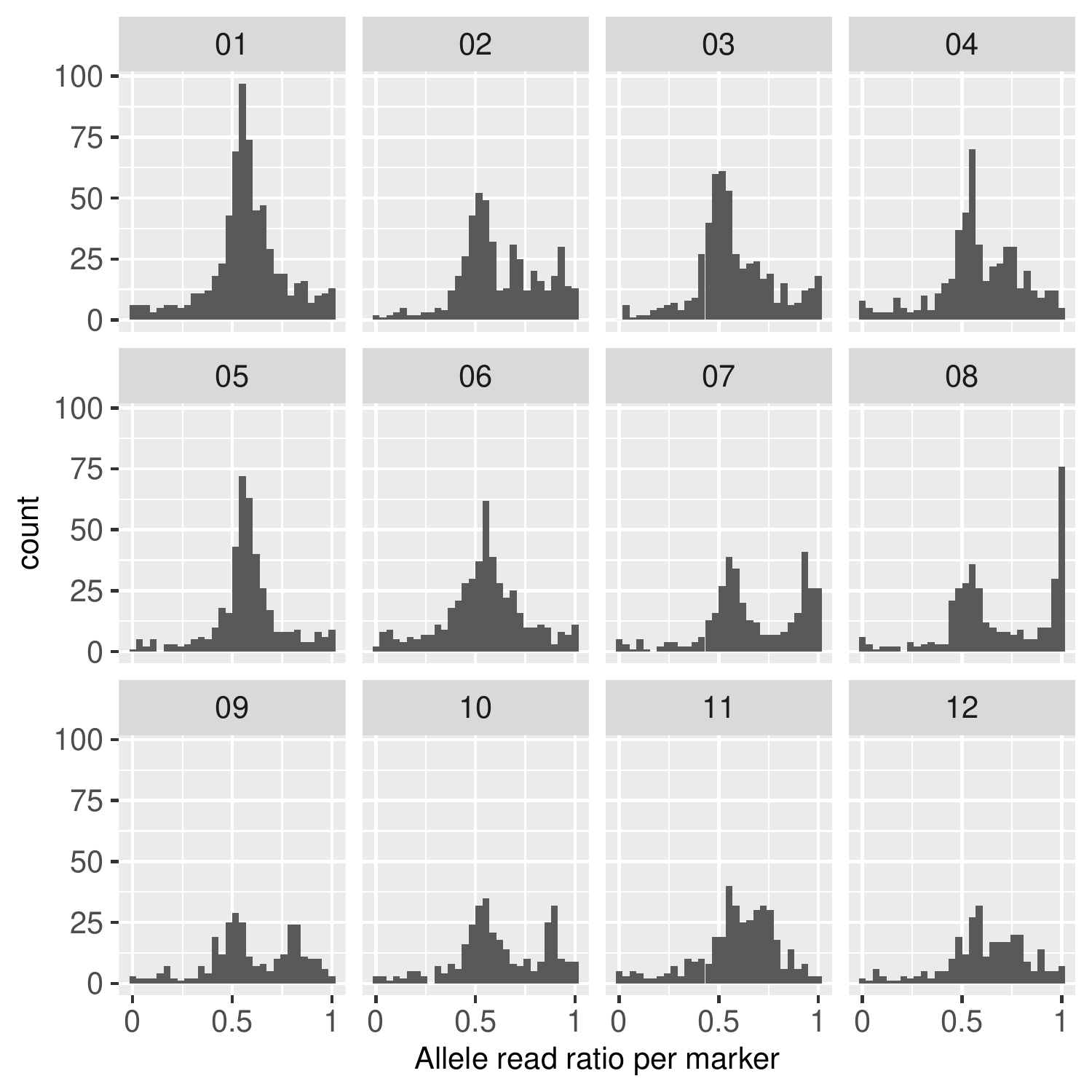
 **Supplemental Figure 1 Allele read ratio per marker observed in the real data**

The histograms of the allele read ratios calculated for each marker are plotted separately for chromosomes 1-12. The allele read ratio greater than 0.5 indicates preferable observation of reference allele reads at the marker. These histograms indicate that many markers tend to have reference reads. The majority of the markers counted in the right-most bin of each histogram were associated with high missing rates, which may lead to apparent allele read biases. The markers in the real data seem to prefer reference allele reads in all chromosomes.

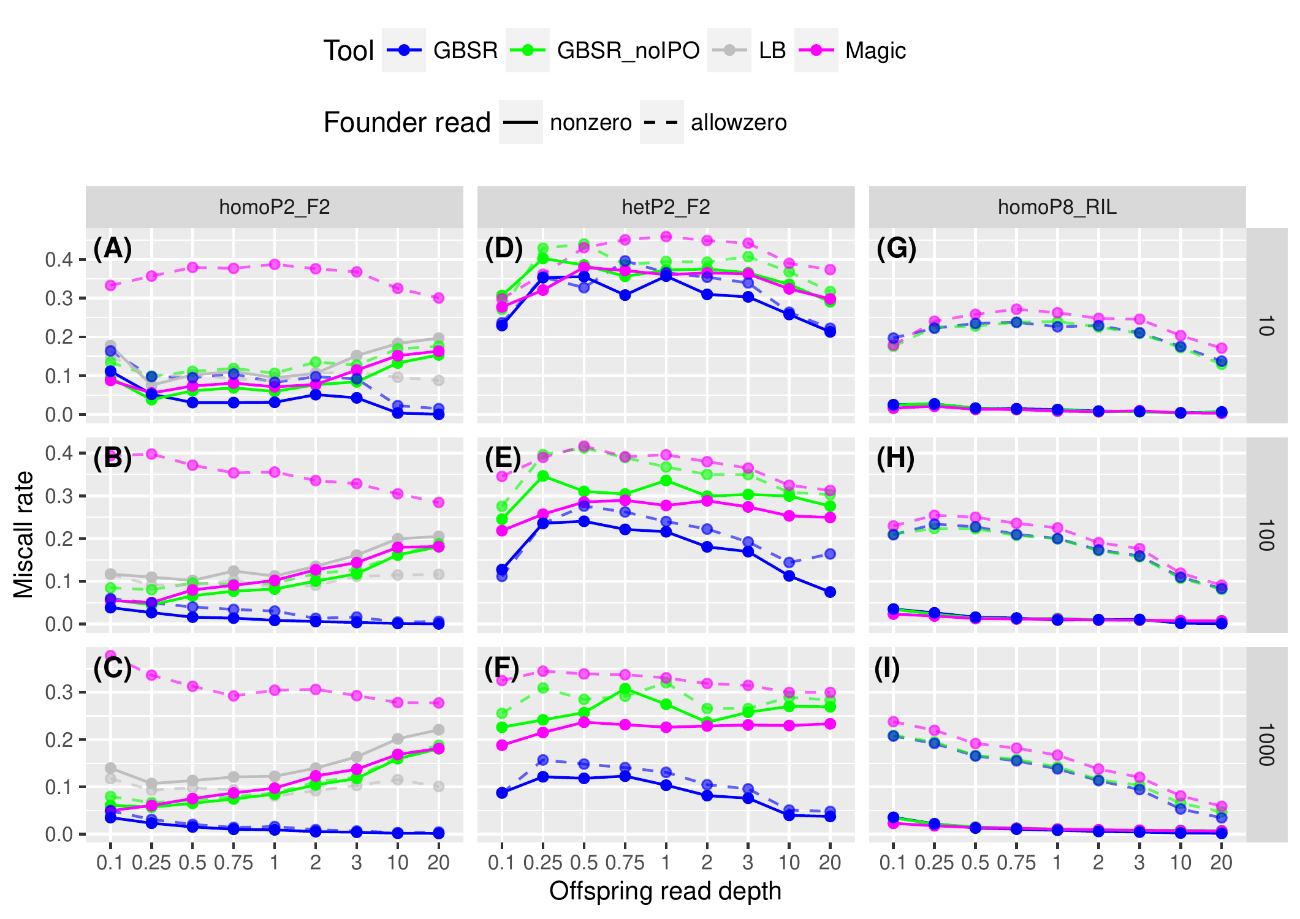

**Supplemental Figure 2 Miscall rates in offspring genotype estimation for the simulation data.**

The plots show miscall rates for the datasets with given read depths for offspring (x axis) in the homoP2_F2 (A-C), hetP2_F2 (D-F), and homoP8_RIL (G-I) scenarios. The rows of the panels indicate the differences in the number of samples in the simulated data as shown in the strips on the right. Solid lines and dashed lines represent the results for the datasets with (allowzero) or without (nonzero) missing of founder reads. “GBSR” and “GBSR_noIPO” represent GBScleanR with and without IPO. “LB” and “Magic” indicate LB-Impute and magicImpute, respectively.

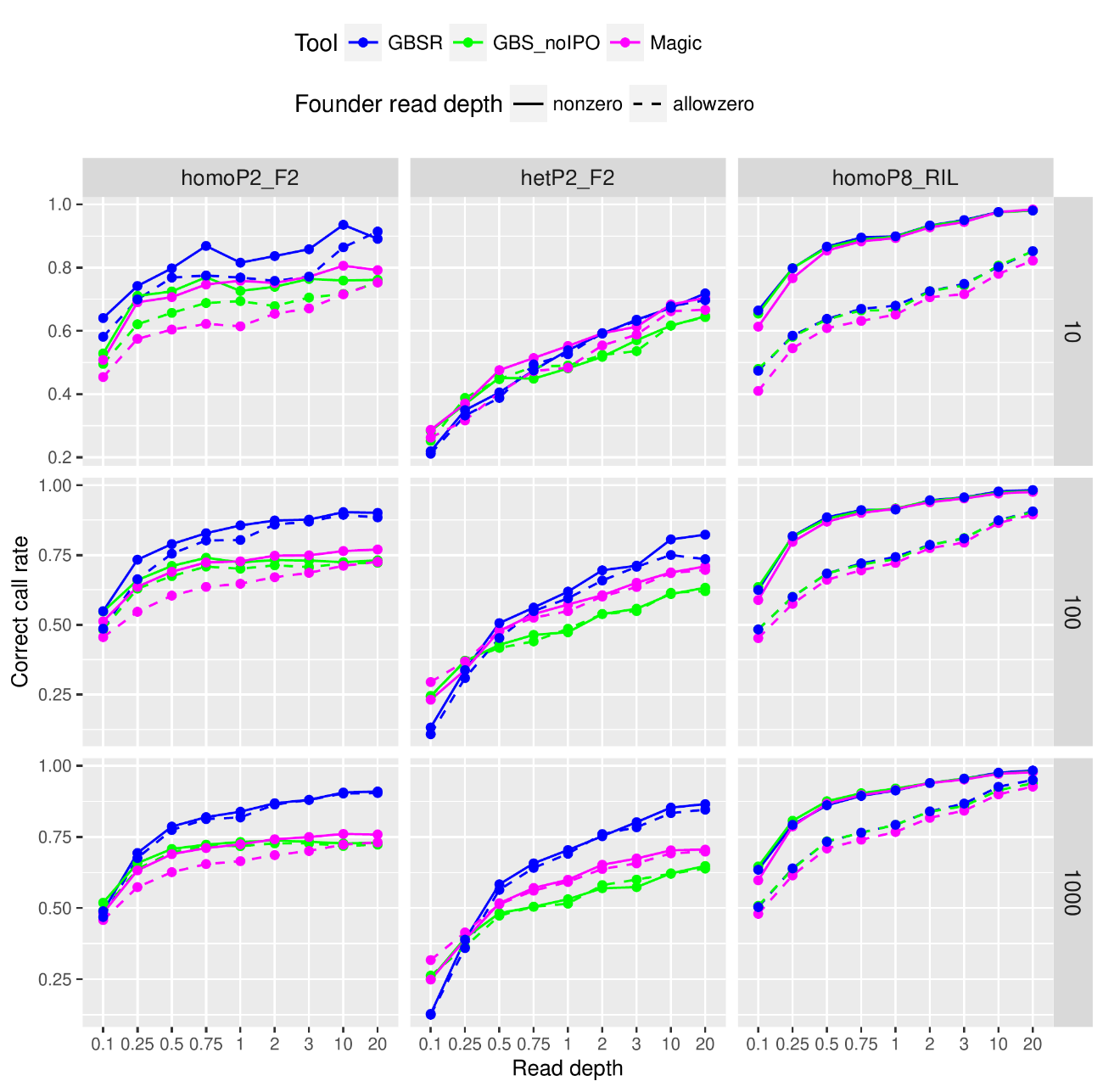

**Supplemental Figure 3 Correct call rates in offspring haplotype estimation for the simulation data.**

The plots show correct call rates of haplotypes for the datasets with given read depths for offspring (x axis) in the homoP2_F2 (A-C), hetP2_F2 (D-F), and homoP8_RIL (G-I) scenarios. The rows of the panels indicate the differences in the number of samples in the simulated data as shown in the strips on the right. Solid lines and dashed lines represent the results for the datasets with (allowzero) or without (nonzero) missing of founder reads. “GBSR” and “GBSR_noIPO” represent GBScleanR with and without IPO. “Magic” indicates magicImpute.

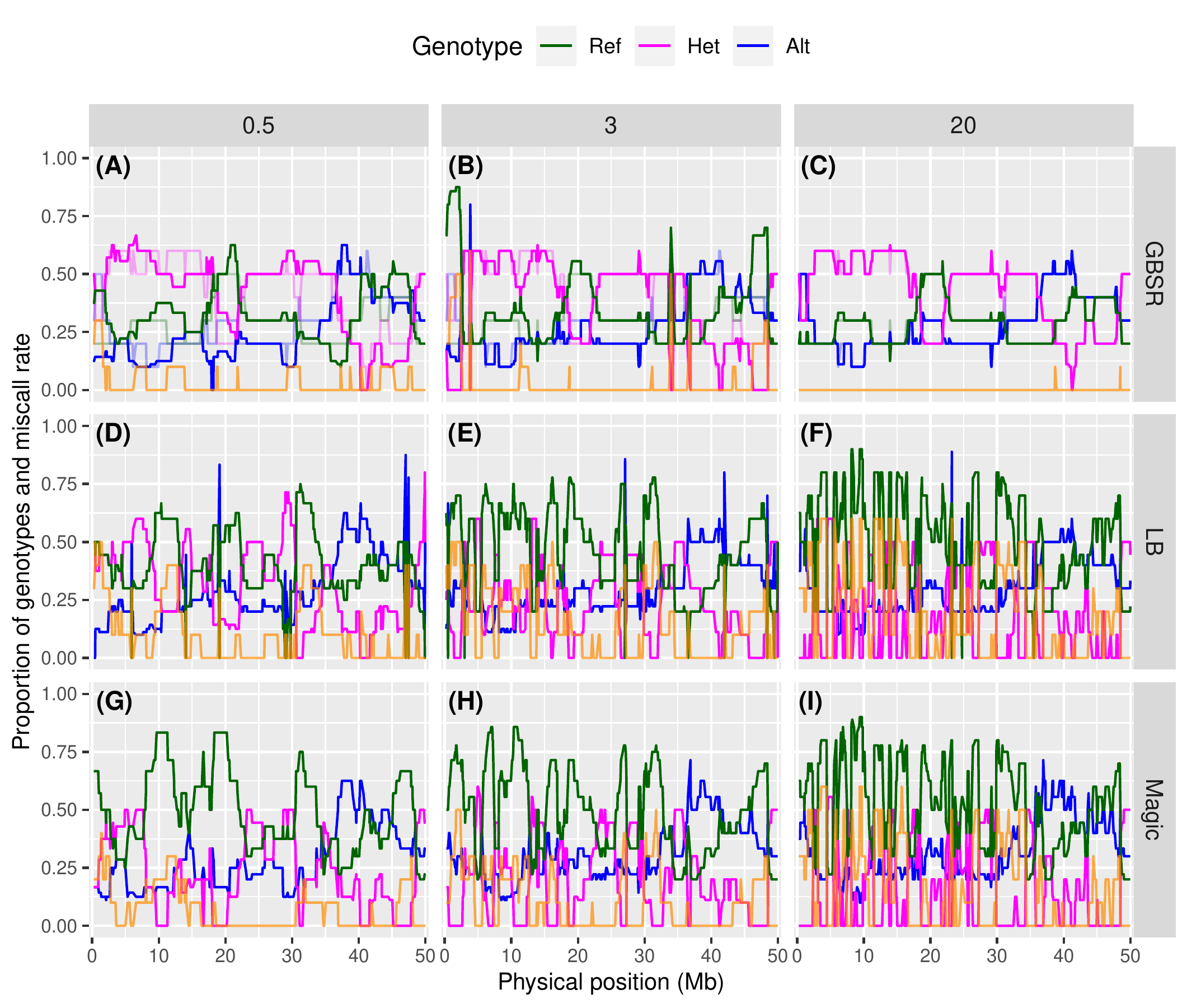

**Supplemental Figure 4 Genotype ratios at markers ordered along a simulated chromosome.** The plots show genotype ratios calculated in the genotype estimation results for the homoP2_F2 dataset with 10 offspring and 620 markers with offspring read depths of 0.5x, 3x, and 20x for offspring and without allowance of no read in founders. ”GBSR” (A-C), ”LB” (D-F), and ”Magic” (G-I) indicates estimated genotype by GBScleanR, LB-Impute, and magicImpute, respectively. Proportions of reference homozygous, heterozygous, and alternative homozygous genotypes are represented by green, magenta, and blue lines, respectively. Orange lines indicate miscall rates at markers. True genotype ratios are indicated by transparent lines in the panels for GBScleanR (A-C).

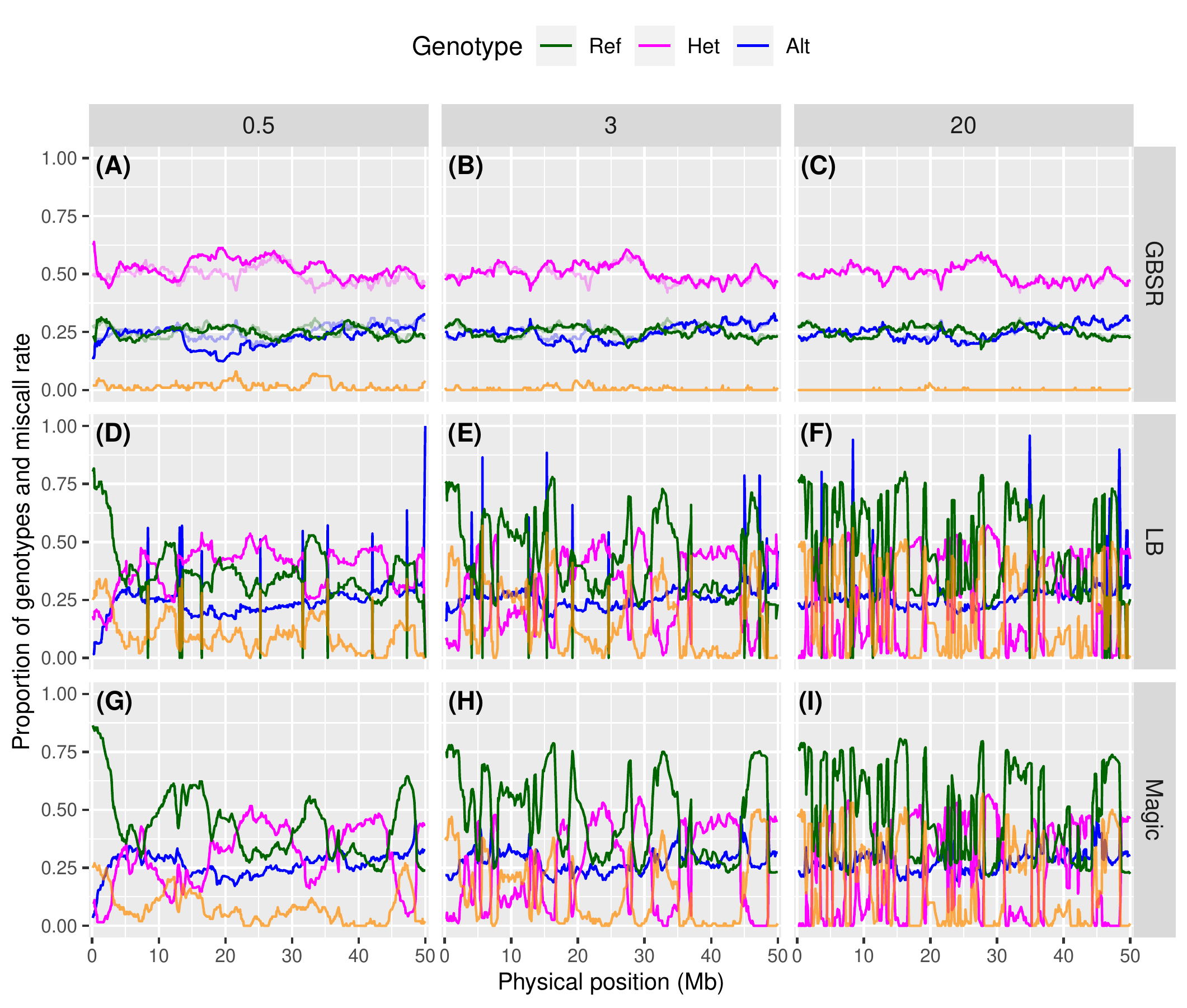

**Supplemental Figure 5** **Genotype ratios at markers ordered along a simulated chromosome.** The plots show genotype ratios calculated in the genotype estimation results for the homoP2_F2 dataset with 100 offspring and 620 markers with offspring read depths of 0.5x, 3x, and 20x for offspring and without allowance of no read. ”GBSR” (A-C), ”LB” (D-F), and ”Magic” (G-I) indicates estimated genotype by GBScleanR, LB-Impute, and magicImpute, respectively. Proportions of reference homozygous, heterozygous, and alternative homozygous genotypes are represented by green, magenta, and blue lines, respectively. Orange lines indicate miscall rates at markers. True genotype ratios are indicated by transparent lines in the panels for GBScleanR (A-C).

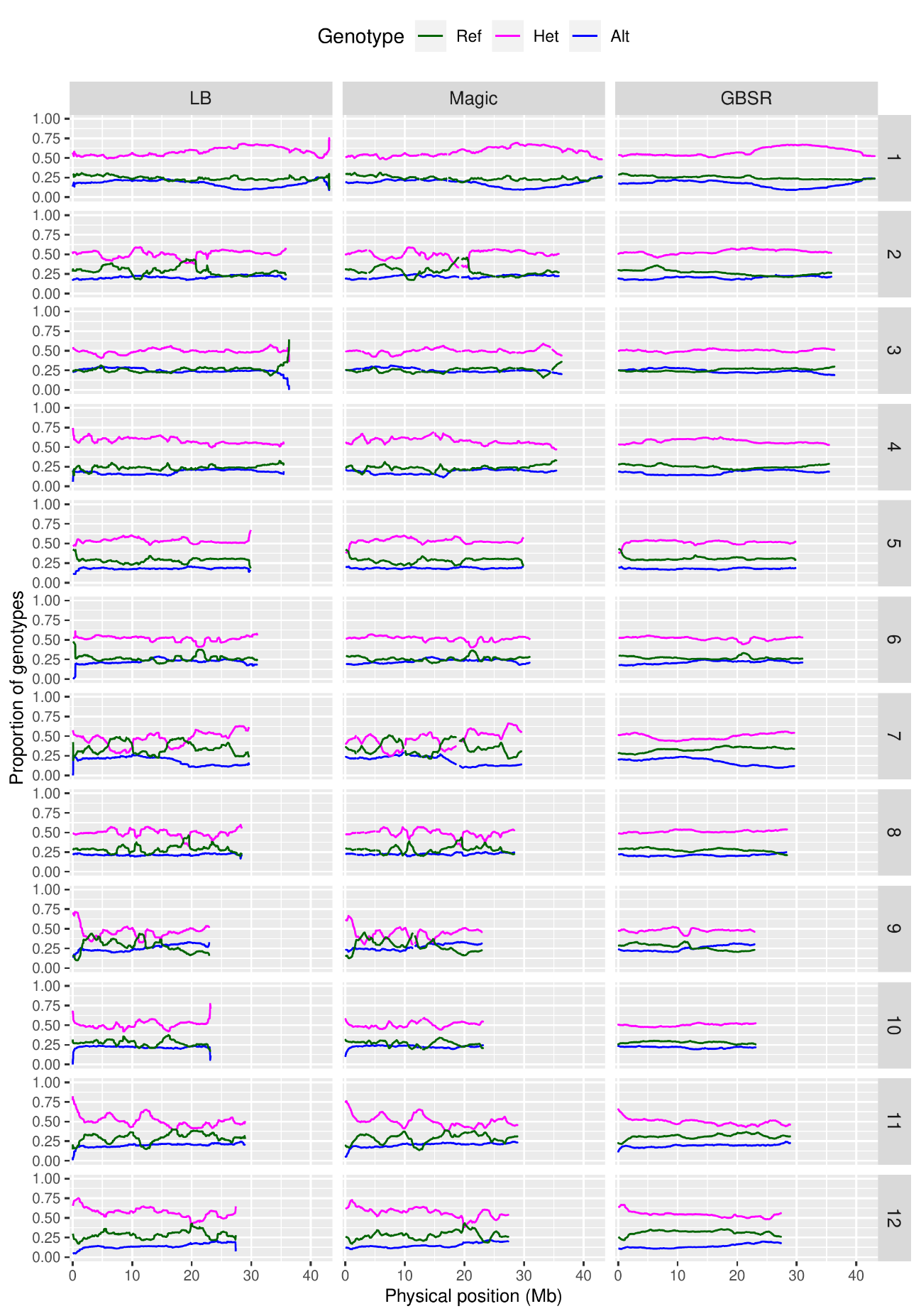

**Supplemental Figure 6 Genotype ratios at markers ordered along chromosomes in the real data.**

“GBSR”, “LB”, and “Magic” indicate GBScleanR, LB-Impute, and magicImpute, respectively. Proportions of reference homozygous, heterozygous, and alternative homozygous genotypes are represented by green, magenta, and blue lines, respectively. Genotype ratios for all chromosomes 12 are shown in this figure

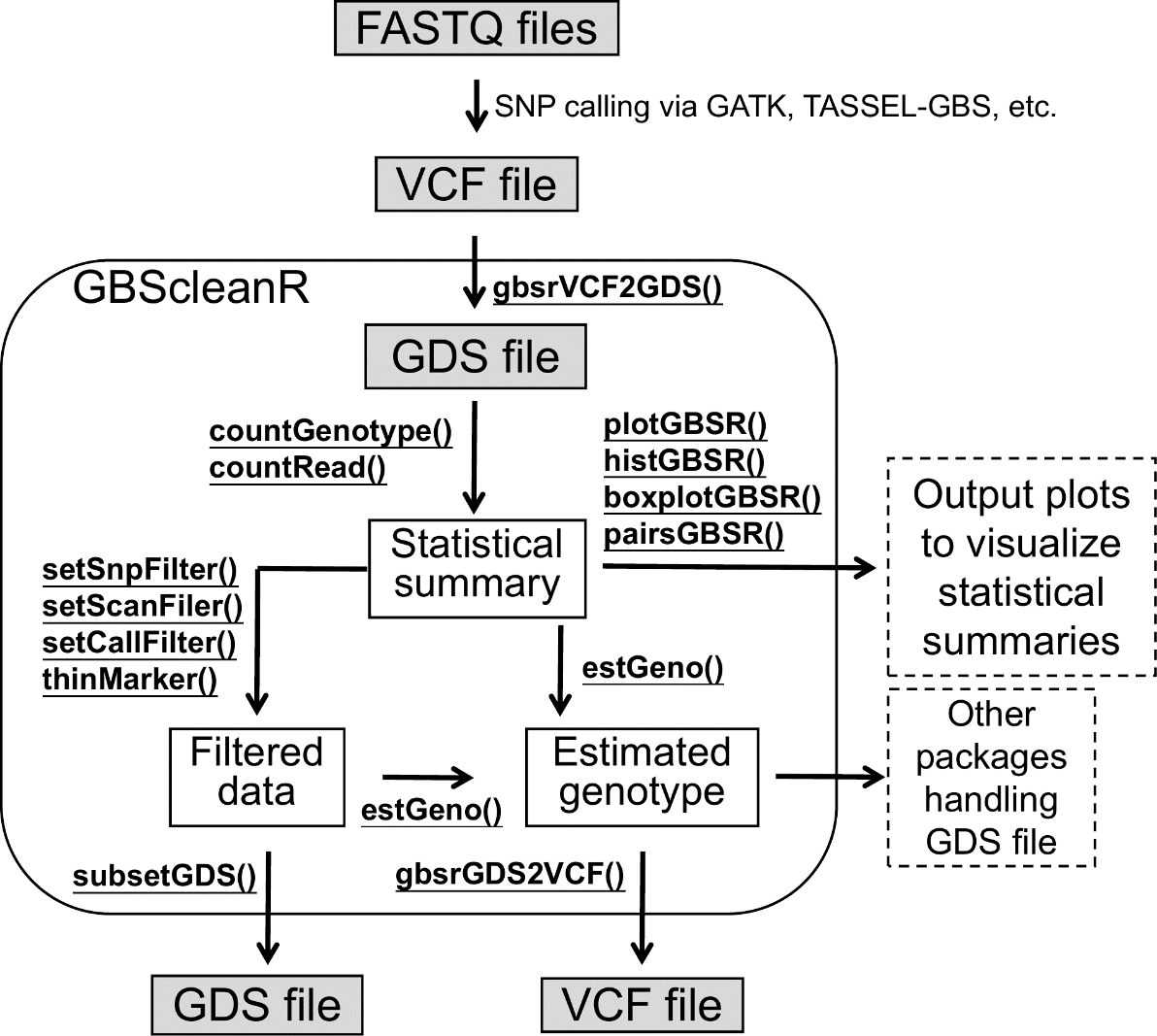

**Supplemental Figure 7 Schematic image of workflow with the GBScleanR package.**

Gray boxes represent data files, while white boxes are data created by GBScleanR in the R environment. Under-lined text indicates the function names of GBScleanR. GBScleanR takes a VCF file generated via SNP callers such as GATK and TASSEL-GBS as inputs. The input VCF file first has to be converted to a GDS file to handle large amounts of genotype data on the R environment with less RAM usage. Statistical summaries of genotype and read counts can be obtained via countGenotype() and countRead(). The summary data can be then visualized using several plotting functions. Users can also filter out samples and markers based on the statistical summaries using setMarFilter() and setSamFilter(). setCallFilter() allow users to filter out single genotype calls at markers based on read counts, while thinMarker() retains one of the markers within a specified stretch in order to reduce the number of markers with redundant information. estGeno()is the core function of GBScleanR that estimates genotype based on given read counts. The filtered genotype data and the estimated genotype data can be output as a new GDS file and a new VCF file by subsetGDS() and gbsrGDS2VCF(), respectively

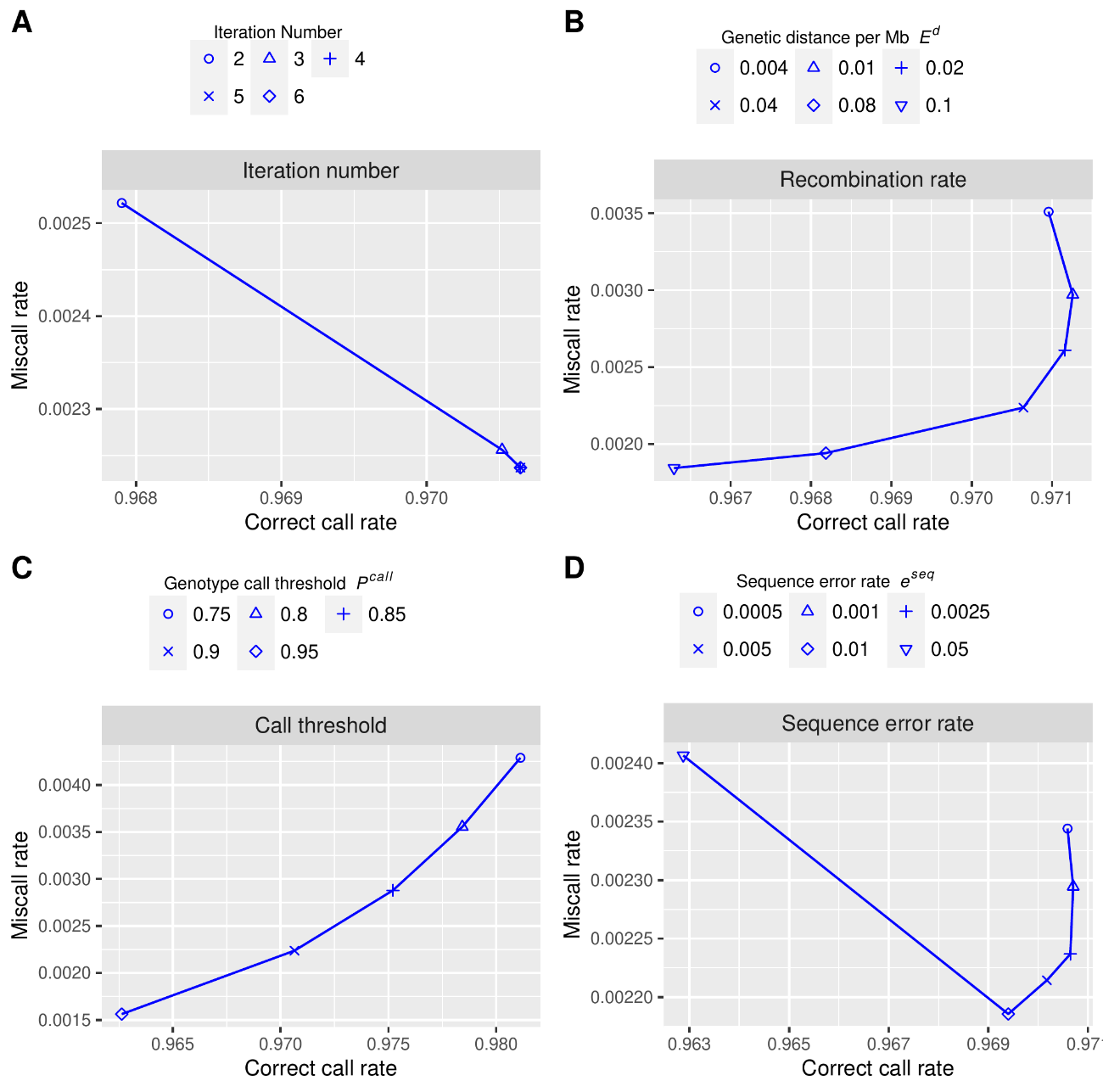

**Supplemental Figure 8 Effects of parameter setting on genotype estimation accuracy.**

Correct call rates and miscall rates at various parameter settings are plotted for iteration number, P_call_ threshold, expected genetic distance per mega base pairs E^d^, and sequence error rate e^seq^ from the top to bottom. Different symbols indicate different settings for each parameter.
